# IL-37 expression reduces acute and chronic neuroinflammation and rescues cognitive impairment in an Alzheimer’s disease mouse model

**DOI:** 10.1101/2021.11.26.470085

**Authors:** Niklas Lonnemann, Shirin Hosseini, Melanie Ohm, Karsten Hiller, Charles A. Dinarello, Martin Korte

**Affiliations:** Department of Cellular Neurobiology, Zoological Institute, TU Braunschweig, 38106 Braunschweig, Germany; Helmholtz Centre for Infection Research, Neuroinflammation and Neurodegeneration Group, 38124 Braunschweig, Germany; Braunschweig Integrated Centre of Systems Biology, TU Braunschweig, Braunschweig, Germany; Department of Computational Biology of Infection Research, Helmholtz Centre for Infection Research, Braunschweig, Germany; Department of Medicine, University of Colorado Denver, Aurora, CO 80045, USA; Department of Medicine, Radboud University; Medical Center, 6525 Nijmegen, The Netherlands

## Abstract

The anti-inflammatory cytokine interleukin-37 (IL-37) is a member of the IL-1 family but not expressed in mice. We used a human IL-37 (hIL-37tg) expressing mouse, which has been subjected to various models of local and systemic inflammation as well as immunological challenges. Those studies demonstrate an immune-modulatory role of IL-37 which can be characterized as an important suppressor of innate immunity. We investigated the functions of IL-37 in the CNS and explored the effects of IL-37 on neuronal architecture and function, microglia phenotype, cytokine production and behavior after inflammatory challenge by intraperitoneal LPS-injection. Reduced spine density, activated microglia phenotype and impaired long-term potentiation (LTP) were observed in wild-type mice after LPS injection, whereas hIL-37tg mice showed no impairment. In addition, we crossed the hIL-37tg mouse with an animal model of Alzheimer’s disease (APP/PS1) to investigate the anti-inflammatory properties of IL-37 under chronic neuroinflammatory conditions. Our results show that IL-37 is able to limit inflammation in the brain after acute inflammatory events and prevent the loss of cognitive abilities in a mouse model of AD.

## Introduction

Neuroinflammation is characterized by glial cell activation and is mediated via pro-inflammatory signals (1, 2). In general, acute inflammation is the initial response of the immune system and is characterized by the rapid production of various cytokines and chemokines, immune cell activation, and phagocytic mechanisms. Although these processes are important to fight pathogens, persistent inflammation can lead to a pathophysiological condition that results in tissue damage and loss of function. It is well known that during inflammation, pro- and anti-inflammatory cytokines interact and affect the outcome. Examples of pro-inflammatory cytokines include interleukin (IL)-1β and TNF-α, both of which elicit a strong acute inflammatory response (3, 4). In contrast, anti-inflammatory cytokines such as IL-4, IL-10 (5, 6) or IL-37 (7) inhibit the action of pro-inflammatory cytokines and hence limit inflammation (8). In the CNS, long-term activation of glial cells leads to neuroinflammation, which is an important hallmark of many neurological disorders, including Alzheimer’s disease (AD), Parkinson’s disease or multiple sclerosis (1). In particular, AD is the most common form of dementia characterized by amyloid plaques and neurofibrillary tangles in the brain tissue. Patients with AD suffer from memory loss, speech disorders, confusion, problems with attention and spatial orientation. Currently, there are no treatment strategies to combat AD despite the considerable clinical need. Innate immune responses and neuroinflammation are thought to contribute significantly to the progression of AD (9).

Therefore, regulatory cytokines that can reduce inflammation in the CNS are of therapeutic value (10, 11). Nevertheless, it remains unclear which of the regulatory mediators act detrimentally or beneficially at each stage of the neuroinflammatory process (1, 12, 13).

For example, the release of IL-1β under pathological conditions can lead to deficits in learning and memory processes as well as in long-term potentiation (LTP) (14–16). In addition, pro-inflammatory cytokines such as TNF-α and IFN-γ have been associated with impairments in both structure and function of hippocampal neurons during viral infection (17). Both IL-1α and IL-1β play a role in autoinflammatory, autoimmune, infectious, and degenerative diseases (18–22). Although IL-1α and IL-1β are encoded by separate genes, both cytokines are structurally related proteins that bind to the type 1 receptor (IL-1R1) and initiate similar innate responses. However, IL-1α is constitutively present in healthy cells but is only released under cell stress conditions, as is the case during inflammation. Moreover, because it is constitutively present, IL-1α acts rapidly to trigger local inflammation. In contrast, IL-1β is not present under healthy state; the IL-1β precursor is not active but requires processing by caspase-1, an intracellular protease, resulting in conversion to an active cytokine that is secreted (23).

In contrast to the IL-1 family members that trigger inflammation, the IL-1 family also includes two members that act as anti-inflammatory signalling agents and are thought to dampen an ongoing innate immune response (7, 24). These are IL-37 and IL-38 (24). For instance, IL-37 serves as a potent mediator to limit the inflammatory responses (25, 26). Therefore, a transgenic mouse model expressing the human splice-variant IL-37b (hIL-37tg) has been generated (7, 25) and showed potent anti-inflammatory and beneficial effects in a wide variety of different pathological conditions affecting diverse organs (7, 27–30). The transgenic mouse was generated to allow continuous expression of IL-37 in almost all cells. This was achieved by using the full-length cDNA of IL-37b through the CMV promoter (7). It is important to note that these IL-37tg mice do not exhibit an abnormal phenotype and reproduce normally (7, 31). IL-37 has been described as a protein that binds to the IL-18 receptor and, unlike IL-18, does not elicit a pro-inflammatory immune response but prevents it and is even involved in an anti- inflammatory response (25). Thus, IL-37 binds the alpha chain of the IL-18 receptor and additionally interacts with IL-1R8 (also described as TIR8 or SIGIRR), rather than recruiting the beta chain of the IL-18 receptor (as is the case with IL-18 binding) (32). This co-localization was also demonstrated in mouse cells (32). Other studies showed the mRNA of IL-18Rα and SIGIRR receptor on astrocytes and microglia (33). In a murine cell line, an important factor of IL-37 expression was also discovered in relation to the instability sequence. Bufler et al. showed that despite strong activation of the CMV promoter, expression of IL-37 was not detected (34). It was also shown that the specific mRNA was rapidly degraded. However, a significant and rapid increase in IL-37 mRNA could be achieved by an LPS stimulus (34). Similarly, administration of recombinant IL- 37 frequently showed beneficial effects on the outcome of various animal models of human diseases (11, 28, 35, 36).

In the present study, we used hIL-37tg mice to investigate the effect of IL-37 expression on the acute inflammatory processes induced by LPS or IL-1β administration. In addition, we crossed the APP/PS1 mouse model of AD with hIL-37tg mice to investigate the role of IL-37 expression during the chronic state of neuroinflammation and the consequences for the progression of AD pathology.

## Results

### Primary microglia from IL-37tg mice exhibit an inflammatory-suppressive response after acute LPS challenge *in vitro*

To investigate the effect of IL-37 on microglial cell activation the release of the pro- inflammatory cytokines IL-6, IL-1β and TNF-α was first analyzed after acute LPS stimulation of WT and IL-37tg microglial cells (Fig. 1A).

**Fig. 1.**
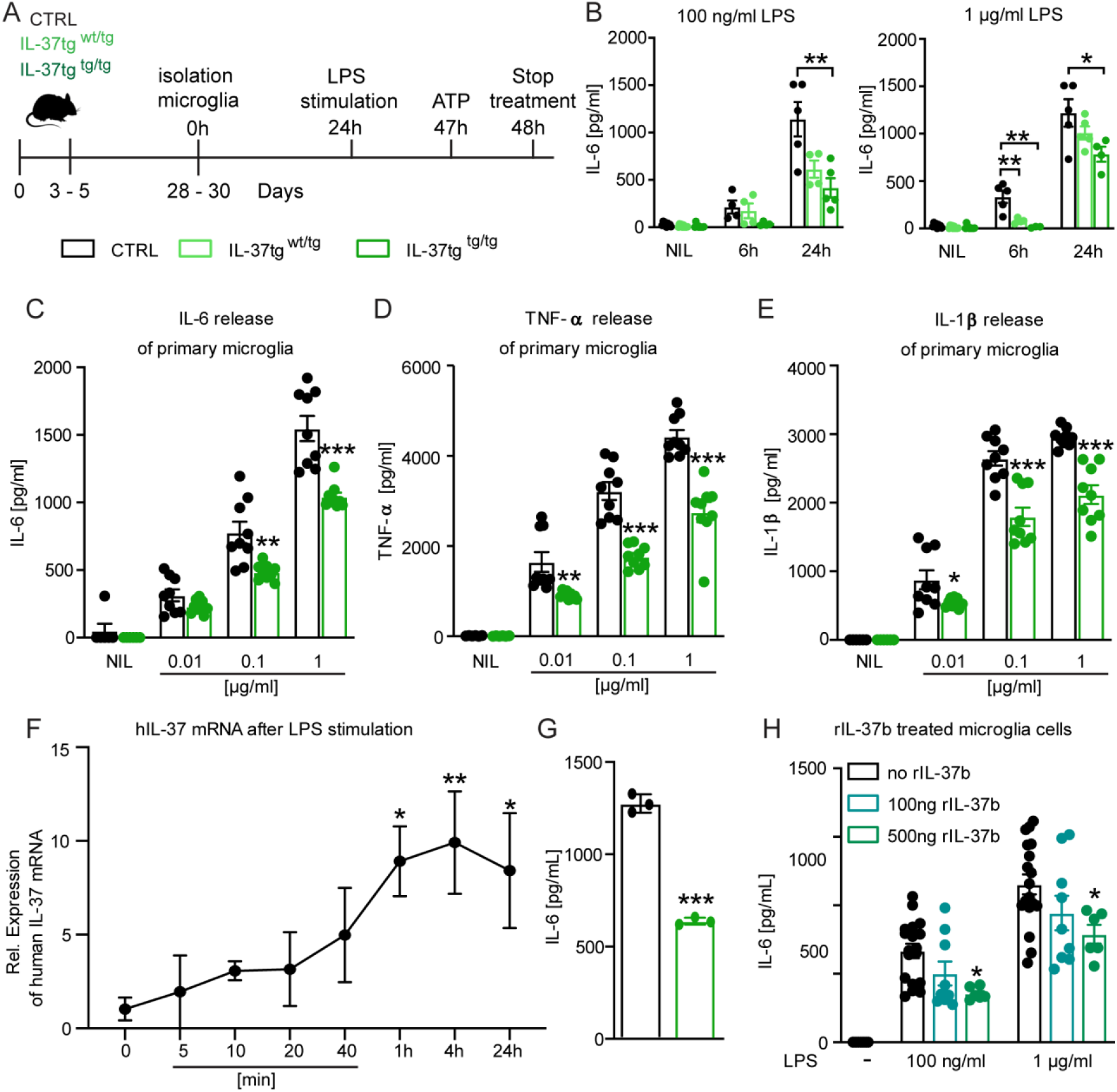
Primary Microglia cells from IL-37tg mice showed reduced pro-inflammatory cytokine release after stimulus of LPS. **(A)** WT and IL-37tg primary microglia cells (P3-5) were plated and stimulated without LPS or with LPS. Cells were treated for 6h **(B)** and 24h **(B-E)**. Cells from homozygote transgenic animals released less pro-inflammatory cytokines IL-6 **(C)**, TNF-α **(D)** and IL-1β **(E)** in different conditions of LPS (1μg/ml - 100ng/ml - 10ng/ml) (B-E n=9). **(F)** *IL37* mRNA was analyzed in IL-37tg microglia cells at defined intervals after the addition of LPS. **(G)** From the same cells of **(F)** IL-6 levels were measured and only detectable after 24h. **(H)** IL-6 levels were measured in WT after addition increasing concentrations of recombinant IL-37b and 100ng LPS (H n=6-18). Data are presented as mean ± SEM. * p < 0.05, ** p < 0.01, *** p < 0.001 compared to WT. (B+F+H: 1-way ANOVA with multiple comparison; C-E+G: t-test).

**Table 1.** Statistical table.

| Figure | Name | Test |  | Multi-comparison |  |  |
| --- | --- | --- | --- | --- | --- | --- |
| 1B | 100ng/ml 6h | 1-way ANOVA | F(2,9)=2,428; P=0,1434 | Tukey's | 0,8947 | WT vs. HET |
|  |  |  |  |  | 0,1460 | WT vs. HOM |
|  | 100ng/ml 24h | 1-way ANOVA | F(2,11)=7,941; P=0,0073 | Tukey's | 0,0539 | WT vs. HET |
|  |  |  |  |  | 0,0067 | WT vs. HOM |
| | 1μg/ml 6h | 1-way ANOVA | F(2,9)=12,74; P=0,0024 | Tukey's | 0,0090 | WT vs. HET |
|  |  |  |  |  | 0,0037 | WT vs. HOM |
| | 1μg/ml 24h | 1-way ANOVA | F(2,11)=3,915; P=0,0520 | Tukey's | 0,3586 | WT vs. HET |
|  |  |  |  |  | 0,0426 | WT vs. HOM |
| 1C | IL-6 10ng/ml | ttest | t=1,742 df=16; P=0,1007 |  |  | WT vs. IL37tg |
|  | IL-6 100ng/ml | ttest | t=3,597 df=16; P=0,0024 |  |  | WT vs. IL37tg |
| | IL-6 1μg/ml | ttest | t=5,127 df=16; P=0,0001 | | | WT vs. IL37tg |
| 1D | TNF-α 10ng/ml | ttest | t=3,376 df=16; P=0,0038 | | | WT vs. IL37tg |
| | TNF-α 100ng/ml | ttest | t=6,774 df=16; P<0,0001 | | | WT vs. IL37tg |
| | TNF-α 1μg/ml | ttest | t=6,264 df=16; P<0,0001 | | | WT vs. IL37tg |
| 1E | IL-1β 10ng/ml | ttest | t=2,390 df=16; P=0,0295 | | | WT vs. IL37tg |
| | IL-1β 100ng/ml | ttest | t=5,027 df=16; P=0,0001 | | | WT vs. IL37tg |
| | IL-1β 1μg/ml | ttest | t=5,852 df=16; P<0,0001 | | | WT vs. IL37tg |
| 1F | IL-37 mRNA | 1-way ANOVA | F(7,15)=2,629; P=0,0550 | Fisher's LSD | 0,0171 | 0h vs. 1h |
|  |  |  |  |  | 0,0086 | 0h vs. 4h |
|  |  |  |  |  | 0,0238 | 0h vs. 24h |
| 1G | IL-6 | ttest | t=20,39 df=4; P<0,0001 |  |  | WT vs. IL37tg |
| 1H | rIL-37 100ng/ml | 1-way ANOVA | F(2,30)=5,135; P=0,0121 | Tukey's | 0,1675 | WT vs. 100ng |
|  |  |  |  |  | 0,0133 | WT vs. 500ng |
| | rIL-37 1μg/ml | 1-way ANOVA | F(2,30)=3,512; P=0,0426 | Tukey's | 0,2457 | WT vs. 100ng |
|  |  |  |  |  | 0,0498 | WT vs. 500ng |
| 2B | Itaconate Treatment | 2-way ANOVA | F(1,20)=174,9;P<0,0001 | Bonferron | <0,0001 | WT NIL vs. LPS |
|  |  |  |  |  | <0,0001 | IL37 NIL vs. LPS |
|  | Itaconate Genotype | 2-way ANOVA | F(1,20)=27,55;P<0,0001 | Bonferron | >0,9999 | NIL WT vs. IL37 |
|  |  |  |  |  | <0,0001 | LPS WT vs. IL37 |
| 2C | Succinate Treatment | 2-way ANOVA | F(1,20)=98,40;P<0,0001 | Bonferron | <0,0001 | WT NIL vs. LPS |
|  |  |  |  |  | 0,0005 | IL37 NIL vs. LPS |
|  | Succinate Genotype | 2-way ANOVA | F(1,20)=17,01;P=0,0005 | Bonferron | >0,9999 | NIL WT vs. IL37 |
|  |  |  |  |  | <0,0001 | LPS WT vs. IL37 |
| 3B | Body weight Treatment | 2-way ANOVA | F(1,17)=220,5;P<0,0001 | Bonferron | <0,0001 | WT NIL vs. LPS |
|  |  |  |  |  | <0,0001 | IL37 NIL vs. LPS |
|  | Body weight Genotype | 2-way ANOVA | F(1,41)=1,486;P=0,2299 | Bonferron | >0,9999 | NIL WT vs. IL37 |
|  |  |  |  |  | 0,0139 | LPS WT vs. IL37 |
| 3D | CD68 Treatment | 2-way ANOVA | F(1,6)=19,52;P=0,0045 | Bonferron | 0,0017 | WT NIL vs. LPS |
|  |  |  |  |  | >0,9999 | IL37 NIL vs. LPS |
|  | CD68 Genotype | 2-way ANOVA | F(1,6)=48,42;P=0,0004 | Bonferron | 0,0167 | NIL WT vs. IL37 |
|  |  |  |  |  | <0,0001 | LPS WT vs. IL37 |
| 3E | IL-1β Treatment | 2-way ANOVA | F(1,7)=9,399;P=0,0182 | Bonferron | 0,0244 | WT NIL vs. LPS |
|  |  |  |  |  | 0,7541 | IL37 NIL vs. LPS |
| | IL-1β Genotype | 2-way ANOVA | F(1,7)=3,182;P=0,0935 | Bonferron | >0,9999 | NIL WT vs. IL37 |
|  |  |  |  |  | 0,0366 | LPS WT vs. IL37 |
| 3F | Iba1 CA1 Treatment | 2-way ANOVA | F(1,25)=5,222;P=0,0311 | Bonferron | 0,0024 | WT NIL vs. LPS |
|  |  |  |  |  | >0,9999 | IL37 NIL vs. LPS |
|  | Iba1 CA1 Genotype | 2-way ANOVA | F(1,34)=4,951;P=0,0328 | Bonferron | >0,9999 | NIL WT vs. IL37 |
|  |  |  |  |  | 0,0027 | LPS WT vs. IL37 |
| 3G | Processes CA1 Treatment | 2-way ANOVA | F(1,46)=1,066;P=0,3073 | Bonferron | >0,9999 | WT NIL vs. LPS |
|  |  |  |  |  | 0,6229 | IL37 NIL vs. LPS |
|  | Processes CA1 Genotype | 2-way ANOVA | F(1,58)=1,238;P=0,2704 | Bonferron | 0,5670 | NIL WT vs. IL37 |
|  |  |  |  |  | >0,9999 | LPS WT vs. IL37 |
| 3H | Iba1 Cortex Treatment | 2-way ANOVA | $F(1,25)=15,97; P=0,0005$ | Bonferroni | 0,0002 | WT NIL vs. LPS |
|  |  |  |  |  | 0,3234 | IL37 NIL vs. LPS |
| | Iba1 Cortex Genotype | 2-way ANOVA | $F(1,34)=5,159; P=0,0296$ | Bonferroni | >0,9999 | NIL WT vs. IL37 |
|  |  |  |  |  | 0,0054 | LPS WT vs. IL37 |
| 3I | Processes Cortex Treatment | 2-way ANOVA | $F(1,104)=9,865; P=0,0022$ | Bonferroni | 0,1401 | WT NIL vs. LPS |
|  |  |  |  |  | 0,0232 | IL37 NIL vs. LPS |
| | Processes Cortex Genotype | 2-way ANOVA | $F(1,104)=0,119; P=0,7308$ | Bonferroni | >0,9999 | NIL WT vs. IL37 |
|  |  |  |  |  | 0,8147 | LPS WT vs. IL37 |
| 4A | LTP WT | 2-way ANOVA | $F(1,28)=4,459; P=0,0438$ | Fisher's LSD<br>Time point 43-80 | <0.05 | WT NIL vs. LPS |
| 4B | LTP IL37 | 2-way ANOVA | $F(1,23)=0,085; P=0,7734$ | | | IL37 NIL vs. LPS |
| 4C | LTP last 5min Treatment | 2-way ANOVA | $F(1,22)=7,887; P=0,0102$ | Bonferroni | 0,0043 | WT NIL vs. LPS |
|  |  |  |  |  | >0,9999 | IL37 NIL vs. LPS |
| | LTP last 5min Genotype | 2-way ANOVA | $F(1,29)=1,552; P=0,2228$ | | | |
| 4D | Spines CA1 Treatment | 2-way ANOVA | $F(1,32)=22,81; P<0,0001$ | Bonferroni | <0,0001 | WT NIL vs. LPS |
|  |  |  |  |  | 0,1145 | IL37 NIL vs. LPS |
| | Spines CA1 Genotype | 2-way ANOVA | $F(1,44)=26,68; P<0,0001$ | Bonferroni | 0,0657 | NIL WT vs. IL37 |
|  |  |  |  |  | <0,0001 | LPS WT vs. IL37 |
| 4E | Spines DG Treatment | 2-way ANOVA | $F(1,26)=3,307; P=0,0805$ | Bonferroni | 0,0062 | WT NIL vs. LPS |
|  |  |  |  |  | >0,9999 | IL37 NIL vs. LPS |
| | Spines DG Genotype | 2-way ANOVA | $F(1,36)=0,529; P=0,4718$ | Bonferroni | 0,5222 | NIL WT vs. IL37 |
|  |  |  |  |  | 0,0295 | LPS WT vs. IL37 |
| 4G | LTP WT | 2-way ANOVA | $F(1,30)=4,925; P=0,0342$ | Fisher's LSD<br>Time point 39-80 | <0.05 | WT NIL vs. LPS |
| 4H | LTP IL37 | 2-way ANOVA | $F(1,26)=0,01395; P=0,9069$ | | | IL37 NIL vs. LPS |
| 4I | LTP last 5min Treatment | 2-way ANOVA | $F(1,27)=4,613; P=0,0409$ | Bonferroni | 0,0285 | WT NIL vs. LPS |
|  |  |  |  |  | >0,9999 | IL37 NIL vs. LPS |
|  | LTP last 5min<br>Genotype | 2-way<br>ANOVA | F(1,29)=1,747;P=0,1966 |  |  |  |
| 5A | Y-Maze Treatment | 2-way<br>ANOVA | F(1,6)=18,25;P=0,0052 | Bonferroni | 0,0046 | WT NIL vs. IL1 |
|  |  |  |  |  | 0,7313 | IL37 NIL vs. IL1 |
|  | Y-Maze pre-treatment | 2-way<br>ANOVA | F(1,6)=9,899;P=0,0199 | Bonferroni | >0,9999 | NIL WT vs. IL37 |
|  |  |  |  |  | 0,0024 | IL1 WT vs. IL37 |
| 5B | IL-6 Treatment | 2-way<br>ANOVA | F(1,16)=264,6;P<0,0001 | Bonferroni | <0,0001 | WT NIL vs. IL1 |
|  |  |  |  |  | <0,0001 | IL37 NIL vs. IL1 |
|  | IL-6 pre-treatment | 2-way<br>ANOVA | F(1,16)=10,27;P=0,0055 | Bonferroni | 0,8327 | NIL WT vs. IL37 |
|  |  |  |  |  | 0,0004 | IL1 WT vs. IL37 |
| 5C | IL-1α Treatment | 2-way<br>ANOVA | F(1,16)=132,0;P<0,0001 | Bonferroni | <0,0001 | WT NIL vs. IL1 |
|  |  |  |  |  | <0,0001 | IL37 NIL vs. IL1 |
| | IL-1α pre-treatment | 2-way<br>ANOVA | F(1,16)=17,47;P=0,0007 | Bonferroni | >0,9999 | NIL WT vs. IL37 |
|  |  |  |  |  | 0,0003 | IL1 WT vs. IL37 |
| 6B | IL-1β | 1-way<br>ANOVA | F(2,19)=5,224;P=0,0156 | Fisher's<br>LSD | 0,0903 | WT vs. APP |
|  |  |  |  |  | 0,2480 | WT vs. APP-IL37 |
|  |  |  |  |  | 0,0046 | APP vs. APP-IL37 |
| 6C | IL-6 | 1-way<br>ANOVA | F(2,21)=3,600;P=0,0452 | Fisher's<br>LSD | 0,0220 | WT vs. APP |
|  |  |  |  |  | 0,5264 | WT vs. APP-IL37 |
|  |  |  |  |  | 0,0466 | APP vs. APP-IL37 |
| 6E | CD68 9-12m | 1-way<br>ANOVA | F(2,11)=14,39;P=0,0008 | Bonferroni | 0,0008 | WT vs. APP |
|  |  |  |  |  | 0,0529 | WT vs. APP-IL37 |
|  |  |  |  |  | 0,04494 | APP vs. APP-IL37 |
| 6F | CD68 6m | 1-way<br>ANOVA | F(2,12)=13,45;P=0,0009 | Bonferroni | 0,0007 | WT vs. APP |
|  |  |  |  |  | 0,0263 | WT vs. APP-IL37 |
|  |  |  |  |  | 0,0985 | APP vs. APP-IL37 |
| 6G | CD68 20-23m | 1-way ANOVA | $F(2,12)=13,38; P=0,0009$ | Bonferroni | 0,0007 | WT vs. APP |
|  |  |  |  |  | 0,0359 | WT vs. APP-IL37 |
|  |  |  |  |  | 0,0383 | APP vs. APP-IL37 |
| 6I | Plaque load Hp | ttest | $t=2,403$ df=74; $P=0,0187$ | | | APP vs. APP-IL37 |
| | Plaque load Cx | ttest | $t=4,953$ df=63; $P<0,0001$ | | | APP vs. APP-IL37 |
| 6J | Plaque size Hp | ttest | $t=2,296$ df=74; $P=0,0245$ | | | APP vs. APP-IL37 |
| | Plaque size Cx | ttest | $t=5,949$ df=63; $P<0,0001$ | | | APP vs. APP-IL37 |
| 6G | Abeta uptake | 1-way ANOVA | $F(2,22)=136; P<0,0001$ | Bonferroni | 0,0001 | WT vs. APP |
|  |  |  |  |  | 0,0001 | WT vs. APP-IL37 |
|  |  |  |  |  | 0,0026 | APP vs. APP-IL37 |
| 7B | Latency WT | 1-way ANOVA | $F(7,88)=5,842; P<0,0001$ | | | WT |
| | Latency APP | 1-way ANOVA | $F(7,72)=3,477; P=0,0029$ | | | APP |
| | Latency APP-IL37 | 1-way ANOVA | $F(7,80)=4,445; P=0,0003$ | | | APP-IL37 |
| | Latency | 2-way ANOVA | $F(2,30)=10,50; P=0,0003$ | Fisher's LSD<br>Day 1 | <0.05 | WT vs. APP |
|  |  |  |  | Day 3-6 | <0.01 |  |
| 7C | PT WT | ttest | $t=11,45$ df=22; $P<0,0001$ | | | NT vs. TQ |
| | PT APP | ttest | $t=0,9874$ df=18; $P=0,3365$ | | | NT vs. TQ |
| | PT APP-IL37 | ttest | $t=3,705$ df=20; $P=0,0014$ | | | NT vs. TQ |
| | PT TQs | 1-way ANOVA | $F(2,30)=8,415; P=0,0013$ | Turkey's | 0,0009 | WT vs. APP |
|  |  |  |  |  | 0,0505 | WT vs. APP-IL37 |
| 7E | LTP | 2-way ANOVA | $F(2,45)=9,286; P=0,0004$ | Fisher's LSD<br>Time point 21-80 | <0.0005 | WT vs. APP |
|  |  |  |  | Fisher's LSD<br>Time point 41-80 | <0.05 | WT vs. APP-IL37 |
| 7F | Mean LTP 20-25min | 1-way ANOVA | $F(2,45)=7,090; P=0,0021$ | Turkey's | 0,0016 | WT vs. APP |
|  |  |  |  |  | 0,5444 | WT vs. APP-IL37 |
| 7G | Mean LTP 75-80min | 1-way ANOVA | $F(2,45)=9,354; P=0,0004$ | Turkey's | 0,0014 | WT vs. APP |
|  |  |  |  |  | 0,0023 | WT vs. APP-IL37 |
| 7I | Spine density | 1-way ANOVA | $F(2,63)=6,318; P=0,0032$ | Turkey's | 0,0026 | WT vs. APP |
|  |  |  |  |  | 0,0536 | WT vs. APP-IL37 |

Data showed that the amounts of IL-6 secreted by microglial cells from homozygous IL- 37 transgenic (IL-37^tg/tg^) mice were significantly reduced compared to microglial cells from control mice after 24 hours of stimulation with 100ng/ml and 1µg/ml LPS and 6 hours after stimulation with 1µg/ml LPS (F(2, 11)=7.941 p=0.007; p=0.007; p=0.054; F(2, 11)=3.915 p=0.052; p=0.043; p=0.359; F(2, 9)=12.74 p=0.002; p=0.004; p=0.009) (Fig. 1B). In addition, microglial cells from heterozygous IL-37tg mice (IL-37^tg/wt^) also showed decreased IL-6 secretion after 6 hours of stimulation with 1µg/ml LPS (F(2, 9)=12.74 p=0.002; p=0.009) (Fig. 1B). Thus, the immunosuppressive effect by the transgenic expression of IL-37 on primary microglia could also be reproduced for the release of IL-6 when other concentrations of the inflammatory stimulus were examined (Fig. 1C) and could be extended by the significantly reduced release of TNF-α (Fig. 1D) and IL-1β (Fig. 1E), compared with primary microglial cells from control mice (IL-6: p=0.1007; p=0.0024; p=0.0001; TNF-α: p=0.0038; p<0.0001; p<0.0001; IL-1β: p=0.0295; p=0.0001; p<0.0001). Interestingly, LPS stimulation caused a detectable increase in the level of transgenic *IL-37* mRNA after 1 hour, which remained elevated in the following hours up to 24 hours after stimulation (F(7, 15)=2.629 p=0.05509; p=0.0171, p=0.0086, p=0.0238) (Fig. 1F). As mentioned above, the amount of IL-6 protein in the supernatant of the same cultures was significantly reduced when IL-37tg microglia was compared to identically stimulated control cells (p<0.0001) (Fig. 1G).

To investigate whether the addition of recombinant IL-37 (rIL-37) protein to microglial cells has an effect on the release of pro-inflammatory cytokines after inflammatory stimulation, primary wild-type microglial cells were pretreated with either 100 ng/mL or 500 ng/mL rIL-37b for 2h and then stimulated with increasing concentrations of LPS. Microglial cells treated with the higher concentration of rIL-37b showed a significant reduction in the release of IL-6 compared to cells treated with rIL-37b vehicle (PBS) (F(2, 30)=5.135 p=0.0121; p=0.0133; F(2, 30)=3.512 p=0.0426; p=0.0498) (Fig. 1H). Taken together, these findings show that microglial cells release lower levels of pro-inflammatory cytokines upon acute inflammatory stimulation in the presence of IL-37.

### Metabolomic profiling of microglial cells from IL-37tg mice reveals an attenuated metabolic response associated with inflammation after LPS stimulation *in vitro*

Recent studies have discovered several metabolic intermediates that contribute directly to immune function. Among them, the TCA cycle-related metabolites succinate and itaconate are novel markers associated with pro-inflammatory macrophage activation and function (37–39).

To address the question of whether microglial cells expressing the anti-inflammatory cytokine IL-37 are further capable of modulating inflammation-induced metabolic changes, we performed intracellular metabolomic analysis of control and IL-37tg microglial cells treated with 10 ng/ml LPS for 24 hours. As expected, many metabolites were significantly increased in control cells treated with LPS, including itaconate and succinate, both of which are metabolic markers of pro-inflammatory activation of macrophages (Fig. 2A) (38, 40). Levels of both metabolites were significantly less elevated in IL-37tg microglial cells, indicating an attenuated pro-inflammatory response at the metabolic level (Fig. 2B and C) (itaconate F(1, 20)=174.9 p<0.0001; p<0.0001; p<0.0001; F(1, 20)=27.55 p<0.0001; p<0.0001; succinate F(1, 20)=98.40 p<0.0001; p<0.0001; p=0.0005; F(1, 20)=17.01 p=0.0005; p<0.0001). This is consistent with the profiles of other metabolites of central carbon metabolism showing modest effects after LPS treatment (Fig. 2A). We conclude that IL-37tg microglial cells release lower levels of pro-inflammatory cytokines after acute LPS induction than control and further that IL-37 expression in microglial cells limits LPS induced pro-inflammatory metabolic reprogramming.

**Fig. 2.**
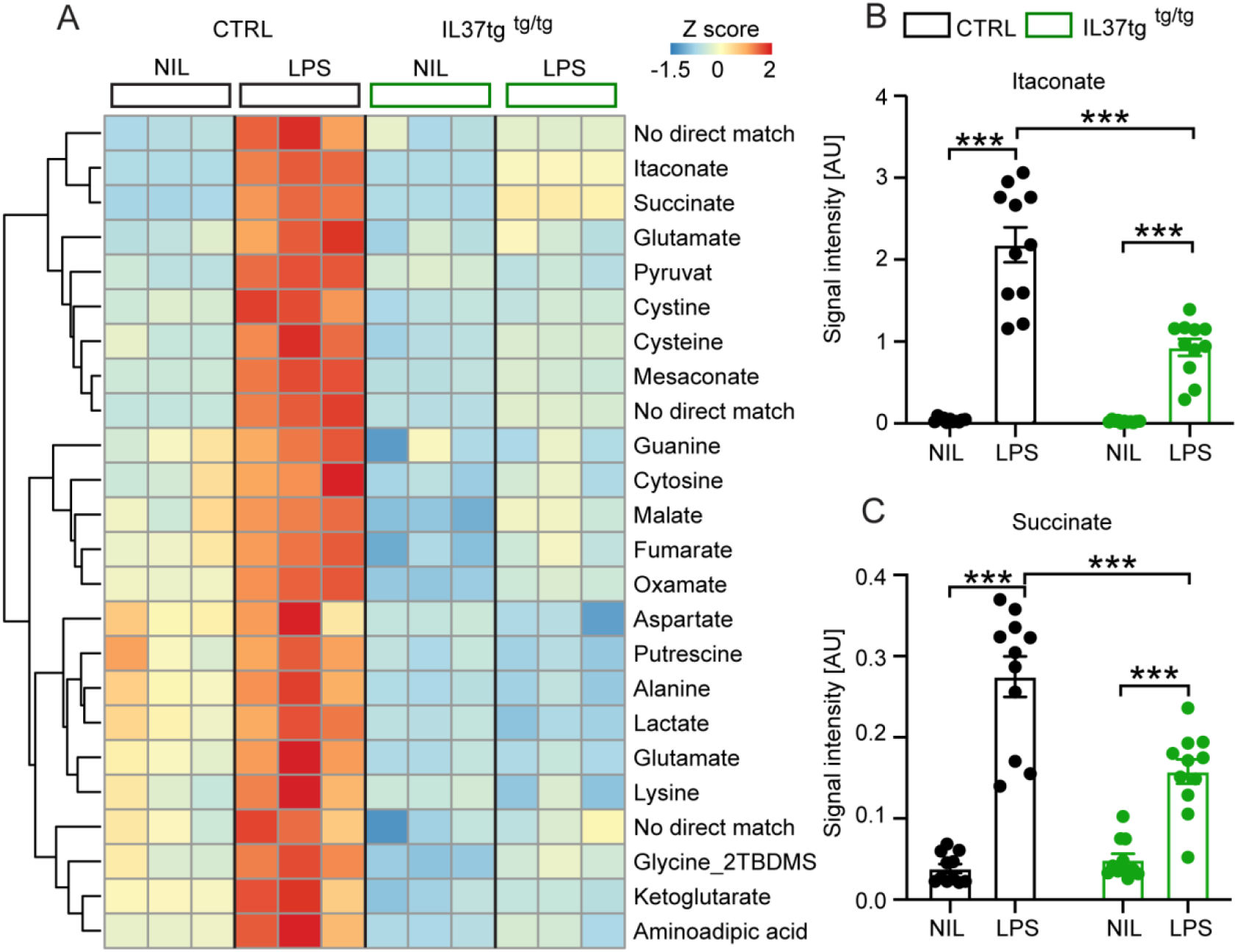
Primary microglia cells isolated from IL-37tg mice showed reduced levels of inflammation-associated intracellular metabolites after stimulus of LPS. WT and IL-37tg microglia were stimulated with 10 ng/ml LPS (due to highly sensitive metabolomics assessments). Metabolomic analysis was performed on these cells. **(A)** Heatmap of identified, significantly changed metabolites upon or without LPS stimulation (described as Z score). **(B-C)** Significant effects of WT treated with LPS was exhibited with respect to Itaconate **(B)** and Succinate **(C)**, whereas these changes were significant reduced for IL-37tg microglia cells treated with LPS compared to WT cells. Data are presented as mean ± SEM. *** p < 0.001 (2-way ANOVA with multiple comparison).

### Microglial activation and inflammatory responses are reduced after *in vivo* LPS challenge in IL-37tg mice

To test whether the anti-inflammatory properties of IL-37 on microglial cells observed in our *in vitro* studies, would also be similar *in vivo*, we injected either saline (vehicle control group) or LPS (2x 0.5 mg/kg) intraperitoneally to adult (3-8 months old) homozygous IL- 37tg (referred to as IL-37tg) mice and age-matched littermate controls (Fig. 3A). We found that both experimental groups showed significant weight loss in response to systemic LPS injection, but the effect was significantly higher in control mice compared to IL-37tg mice (weight loss F(1, 17)=220.5 p<0.0001; p<0.0001; p<0.0001; F(1, 41)=1.486 p=0.2299; p=0.0139) (Fig. 3B). Next, we examined the effect of systemic administration of LPS on neuroinflammation by using brain homogenates for ELISA assays and further isolated microglia from the brains of controls and LPS-treated animals for FACS analysis. We observed that control mice treated with LPS had a significantly higher percentage of CD68-expressing microglial cells (gated on the CD11b^+^/CD45_low_ population) (CD68 F(1, 6)=19.52 p=0.0045; p=0.0017; p>0.9999; F(1, 6)=48.42 p=0.0004; p=0.0001) (Fig. 3C-D) compared to cells obtained from saline treated control animals. Likewise, IL-1β levels in the CNS of LPS-treated WT mice were significantly higher than those of untreated WT controls (IL-1β F(1, 7)=9.399 p=0.0182; p=0.0244; p=0.7541; F(1, 7)=3.182 p=0.0935; p=0.0366) (Fig. 3E). In contrast, IL-37tg animals showed no significant change when challenged with LPS in the same manner (Fig. 3C-E).

**Fig. 3.**
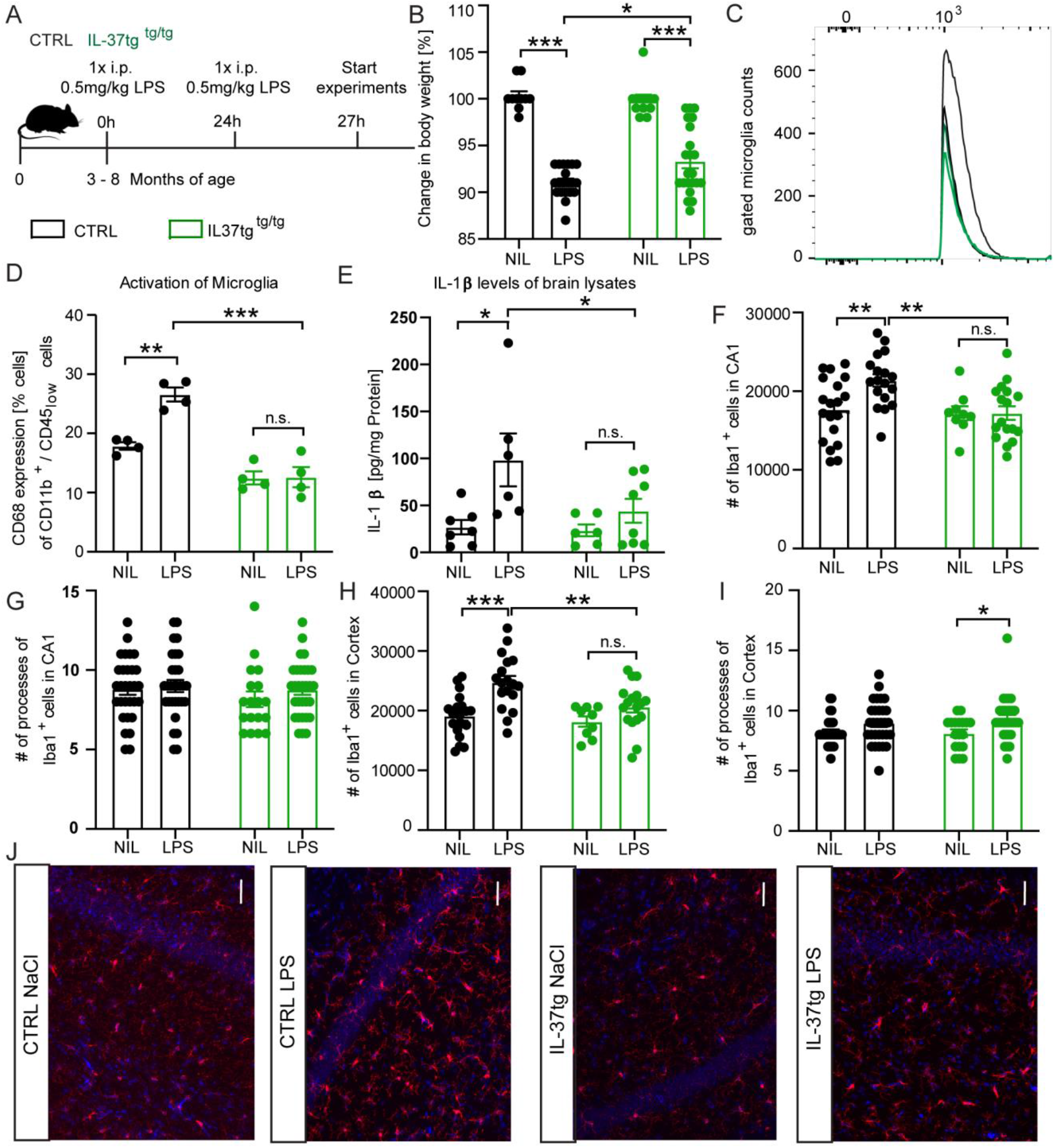
IL-37tg mice showed reduced pro-inflammatory cytokine release and less activated microglia after stimulus of LPS. **(A)** WT and IL-37tg mice were stimulated with saline or LPS. **(B)** IL-37tg animals exhibited a significant lower loss of weight compared to WT mice. However, also IL-37tg mice had a significant change in weight compared to saline treated mice. The activation of microglia cells was analyzed by using the FACS method. **(C-D)** Microglia cells were identified as CD11b+ and CD45low cells and analyzed for their CD68 expression. IL-37tg mice showed a reduced percentage of cells with CD68 expression compared to WT mice after LPS stimulation **(C-D)**. **(E)** In addition, WT mice represented a significant elevation of IL-1β levels after treatment with LPS, whereas IL-37tg mice did not. **(F-I)** The morphological analysis of microglia cells showed an increased number of IBA-1-positive cells in WT animals treated with LPS compared to saline treated ones. In contrast there is no enhanced IBA-1-positive cell numbers in IL-37tg animals after LPS stimulation. **(J)** Representative images of IBA-1 positive cells (red) and DAPI (blue); scale bar 40 µm. Data are presented as mean ± SEM. * p < 0.05, ** p < 0.01, *** p < 0.001, (B-I: 2-way ANOVA with multiple comparison).

An important activation hallmark of microglial cells is reflected by the total number of microglial cells in the brain parenchyma and the number of primary processes of these CNS resident cells. For example, the higher number of IBA-1-positive cells and a reduced number of primary processes correlate with increased microglial activation (41–43). Therefore, we performed immunostaining with the known microglial marker IBA-1 on tissue sections obtained from the brains of animals of both genotypes treated with either saline or LPS. A significant increase in the number of IBA-1 positive cells was observed in the hippocampal CA1 subregion and in the cortex of LPS treated animals compared with saline-treated control animals. However, in the IL-37tg mice, LPS failed to significantly increase microglial numbers (IBA1 CA1 F(1, 25)=5.222 p=0.0311; p=0.0024; p>0.9999; F(1, 34)=4.951 p=0.0328; p=0.0027;; IBA1 Cx F(1, 25)=15.97 p=0.0005; p=0.0002; p=0.3234; F(1, 34)=5.159 p=0.0296; p=0.0054) (Fig. 3F and H). Further analysis of the number of microglial primary processes showed no differences between animals in the control and IL-37tg group treated with either saline or LPS, except that microglial primary processes in the cortex of IL-37tg mice treated with LPS were significantly increased compared with saline-treated control mice which may indicate more branched microglial characteristics (processes CA1 F(1, 46)=1.066; p>0.9999; p=0.6229; processes Cx F(1, 104)=9.865; p=0.1401; p=0.0232) (Fig. 3G and I). In summary, these findings demonstrate the anti-inflammatory effects on the brain induced by transgenic expression of IL-37 in IL-37tg mice.

### IL-37tg mice are protected from functional and structural neuronal deficits after LPS challenge

Neuroinflammation has previously been shown to affect hippocampal network function (17, 44). Because the present data indicate a significant reduction of neuroinflammation after a systemic LPS challenge, we hypothesized that IL-37 might also have a beneficial effect on hippocampal network function and structure. Therefore, as described above, we injected control and IL-37tg animals with either saline or LPS for subsequent analysis of neuronal function and structure. First, we examined long-term synaptic plasticity, the ability of synapses to change their transmission strength, which is considered a cellular correlate of learning and memory processes (45). For this purpose, we induced long-term potentiation (LTP) at the Schaffer collateral CA3 to CA1 pathway in the hippocampus. After 20 minutes of baseline recording, we observed a significantly impaired LTP in the acute hippocampal slices of control mice treated with LPS compared to saline (F(1, 28)=4.459 p=0.0438) (Fig. 4A). In contrast, IL-37tg animals treated with LPS did not show comparable deficits in synaptic plasticity when compared to saline-treated IL-37tg animals (F(1, 23)=0.0849 p=0.7734) (Fig. 4B). These differences were also evident in the maintenance phase of LTP (last 5 minutes of the measurement) (last 5min F(1, 22)=7.887 p=0.0102; p=0.0043; p>0.9999) (Fig. 4C).

**Fig. 4.**
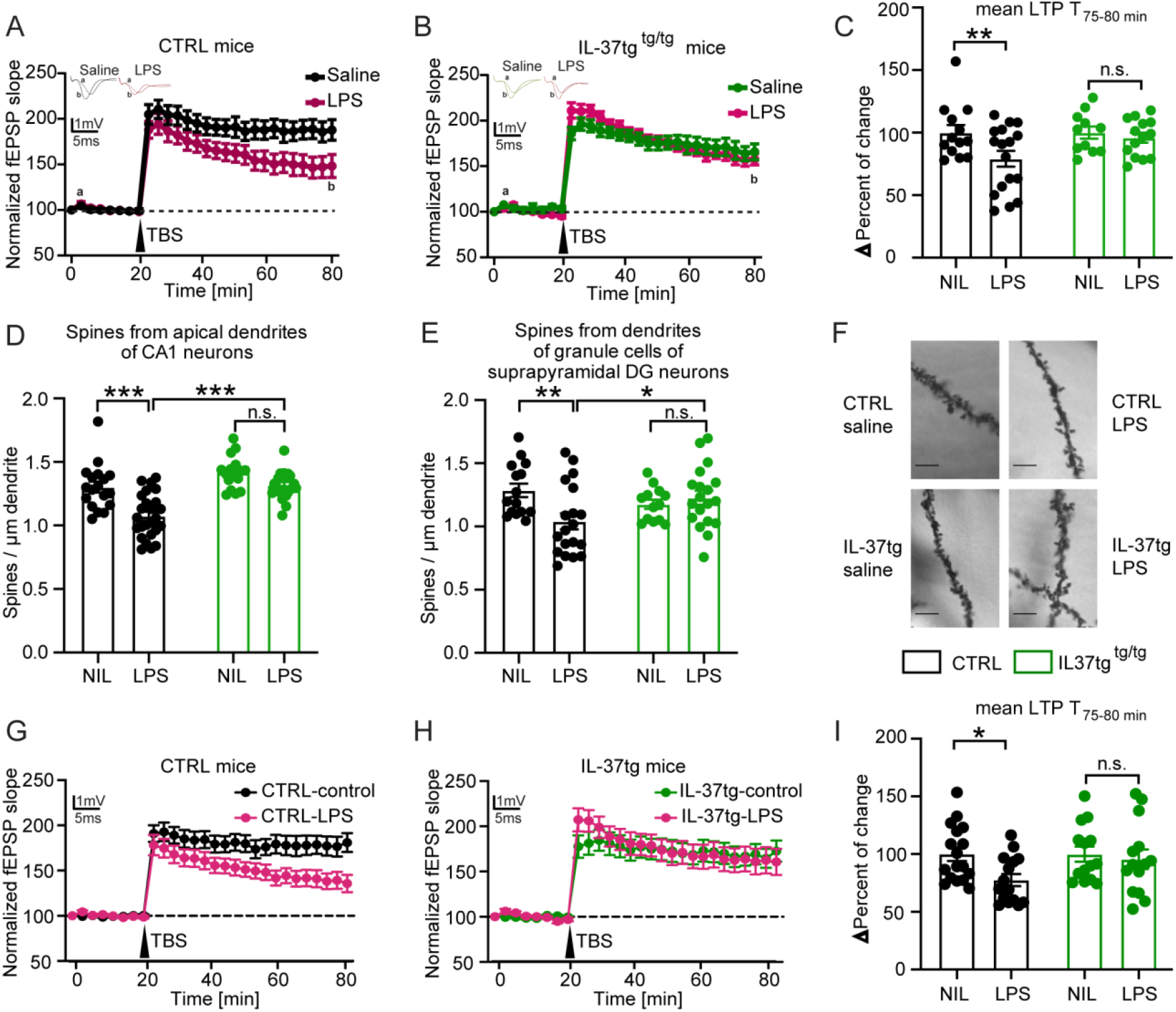
IL-37tg mice showed a rescued synaptic plasticity and a restored spine density loss after stimulus of LPS compared to WT animals. **(A)** WT animals stimulated with LPS showed a significant impairment in LTP induced by theta-burst stimulation (TBS) compared to WT treated with saline. **(B)** IL-37tg mice, on the other hand showed no significant impairment in LTP after LPS treatment. **(C)** The mean LTP attitude (average of 55-60min after TBS) was significant lower in WT mice treated with LPS, whereas IL-37tg mice showed no significant differences. **(D-F)** Dendritic spine density in apical neurons of CA1 hippocampal neurons and in the area of DG was significantly diminished in WT mice treated with LPS, whereas the spine density of IL-37tg animals treated with LPS was not affected **(D and E**). **(F**) Representative images of dendritic spines of hippocampal CA1 neurons in the tested groups were presented; scale bar 5 µm. **G)** WT acute slices stimulated with LPS showed a significant impairment in LTP induced by theta-burst stimulation (TBS) compared to WT acute slices treated with ACSF. **(H)** In contrast, acute slices from IL-37tg mice showed no significant impairment in LTP after LPS treatment. **(I)** The mean LTP attitude (average of 55-60min after TBS) was significant lower in acute slices from WT mice treated with LPS, whereas slices from IL-37tg mice showed no significant differences. Data are presented as mean ± SEM. * p < 0.05, *** p < 0.001, (A-I: 2-way ANOVA with multiple comparison).

To assess the impact of IL-37 on neuronal structure, we analyzed the dendritic spine density of hippocampal neurons from all experimental groups. Dendritic spines are small protrusions representing the postsynaptic part of excitatory synapses and were counted on the apical dendrites of CA1 neurons as well as on the dendrites of dentate gyrus neurons. We found a significant decrease in the density of dendritic spines of CA1 pyramidal neurons and dentate gyrus cells in control mice treated with LPS compared to those treated with saline (spines CA1 F(1, 32)=22.81 p<0.0001; p<0.0001; p=0.1145; F(1, 44)=26.68 p<0.0001; p<0.0001; spines DG F(1, 26)=3.307 p=0.0805; p=0.0062; p>0.9999; F(1, 36)=0.529 p=0.4718; p=0.0295) (Fig. 4D-F). Similar to the LTP data, the density of dendritic spines of neurons from IL-37tg animals was unchanged when LPS- and saline-injected mice were evaluated (Fig. 4D-F). In conclusion, these findings demonstrate a negative effect of systemic administrated LPS on synaptic plasticity that can be restored by IL-37.

Because peripheral cells in IL-37tg mice also produce IL-37 after LPS challenge, the question arose whether IL-37 would have a similar protective effect in the CNS upon LPS stimulation. To accurately demonstrate the beneficial effect of IL-37 in the CNS without the influence of peripheral cells producing IL-37, another electrophysiological experiment was performed. Here, acute slices from control and IL-37tg animals were prepared and stimulated with LPS (10 µg/ml) in ACSF for 2 hours after a resting period before recording. In the control groups, the sections were placed in ACSF only. These further results showed that direct LPS stimulation of acute slices from IL-37tg mice appeared to be sufficient to reverse the impairments in LTP after LPS administration. This is because in contrast to no significant differences in LTP with or without LPS in IL- 37tg acute slices (LTP WT: F(1, 30)=4.925 p 0.05) (Fig. 4G), a significant reduction in LTP was observed in control acute slices when treated with LPS (LTP IL-37tg: F(1, 26)=0.0139 p=0.9069) (Fig. 4H). This effect was also evident in the data from the last 5 minutes of the measurement, which represents the maintenance phase of LTP (last 5min F(1, 27)=4.613 p=0.0409; p=0.0285; p 0.9999) (Fig. 4I). These data clearly demonstrate that local IL-37 in the brain can prevent the deleterious effects induced by LPS for brain function.

### Recombinant IL-37 reduces inflammatory response *in vivo* and relieves short-term memory impairment induced by pro-inflammatory cytokine stimulation

To investigate the anti-inflammatory properties of the recombinant IL-37 (rIL-37) protein *in vivo*, wild-type mice were pretreated for 3 consecutive days with either 300ng of rIL-37 per animal (i.p.) or an equivalent amount of vehicle (saline, control group). On day 4, the animals were injected with 60ng i.p. IL-1β or saline as control (Figure 5A). After another 24 hours, the animals were trained and tested with the Y-Maze behavioral test to assess short-term memory via the mouse’s natural willingness to explore a new territory. The score for spontaneous alternation depends on the mouse tendency to seek out a less recently entered arm of the maze. Hence, this test also measures spatial, hippocampus dependent memory function.

**Fig. 5.**
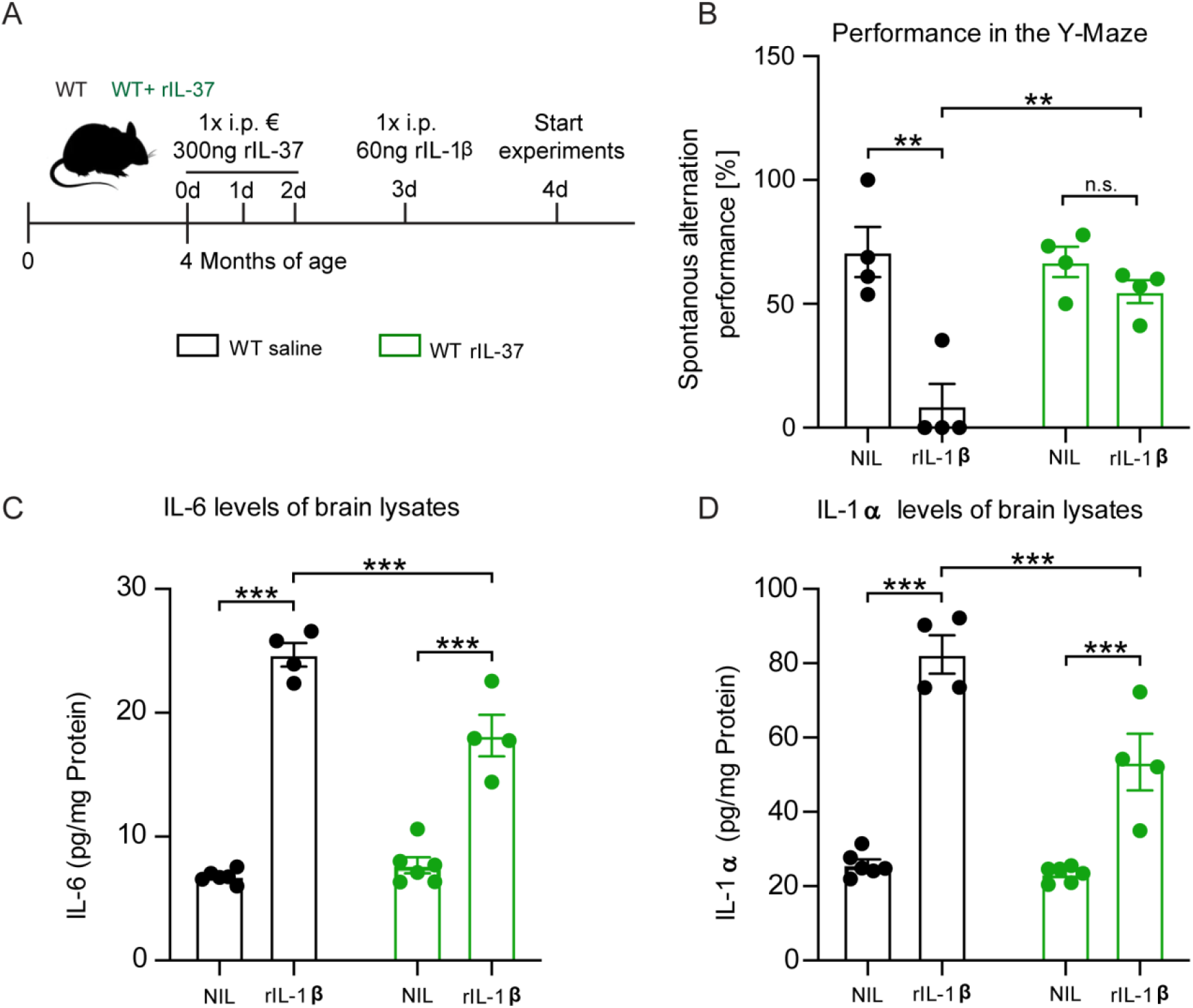
Injection of recombinant IL-37 in WT mice showed restored cognitive deficits and reduced pro-inflammatory cytokine release after inflammatory stimulation of IL-1β. **(A)** WT mice were pre-treated with either saline or rIL-37 on three consecutive days stimulated with saline or IL-1β. **(B)** WT mice treated with saline and IL-1β failed in the performance of the Y-maze test whereas WT mice treated with rIL-37 and IL-1β performed the test without deficits. **(C and D)** Although pro-inflammatory cytokine levels of IL-6 and IL-1a were significantly increased in WT mice treated with rIL-37 compared to the control group, rIL-37 treated mice showed a significant reduction in cytokine levels compared to saline treated after immune stimulation with IL-1β. Data are presented as mean ± SEM. ** p < 0.01, *** p < 0.001, (B-D: 2-way ANOVA with multiple comparison).

Without rIL-37 pretreatment, we detected a significant performance deterioration in IL- 1β-injected mice compared to the corresponding saline control group (Y-Maze F(1, 6)=18.25 p=0.0052; p=0.0046; p=0.7313) (Fig. 5B). In contrast, pretreatment with rIL-37 protected the mice from behavioral deficits after IL-1β (F(1, 6)=9.899 p=0.0199; p=0.0024). To analyze the inflammatory mediators in the CNS of these animals, the levels of the pro-inflammatory cytokines IL-6 and IL-1α were measured in brain lysates. We observed that animals stimulated with rIL-1β always showed significantly increased levels of both cytokines when compared to the corresponding controls (IL-6 F(1, 16)=264.6 p<0.0001; p<0.0001; p<0.0001; F(1, 16)=10.27 p=0.0055; p=0.0004; IL-1α F(1, 16)=132.0 p<0.0001; p<0.0001; p<0.0001) (Fig. 5C-D). However, rIL-37-pretreatment resulted in significantly lower levels of IL-6 and IL-1α in the brains of rIL-1β immune stimulated mice (F(1, 16)=17.47 p=0.0007; p=0.0003) (Fig. 5C-D). Taken together, these data indicate the beneficial effects of IL-37 on cognition and synaptic function in inflammatory stimulated mice.

### IL-37 shows beneficial effects on neuronal deficits and microglia activation in APP/PS1-IL37tg animals

Our findings demonstrated the protective properties of IL-37 on microglial activation, pro-inflammatory mediator production, impaired cognition, and defect long-term potentiation (LTP) after inflammatory challenge. In order to investigate the ability of IL-37 to reduce chronic inflammation, we next analyzed transgenic APP/PS1 mice, which serve as a widely used animal model for Alzheimer’s disease (46), and crossed this mouse strain with hIL-37tg animals (7). First, we examined the levels of pro-inflammatory cytokines in the brains of 9-12 months old control, APP/PS1and APP/PS1-IL37tg transgenic mice (Fig. 6B-C). Compared to control, APP/PS1 animals exhibited higher amounts of IL-6 and IL-1β, while APP/PS1-IL37tg mice showed no increase in these pro-inflammatory cytokines levels (IL-1β F(2, 19)=5.224 p=0.0156; p=0.0903; p=0.248; p=0.0046; IL-6 F(2, 21)=3.6 p=0.0452; p= 0.022; p=0.5264; p=0.0466) (Fig. 6B-C). We then examined microglial cell activation by FACS analysis. For this purpose, the percentage of cells expressing the activation marker CD68 (47–49) by microglia (identified as a CD11b^+^ / CD45_low_ cell population of the brain) over time was analyzed (Fig. 6D-G). In the brains of 9-12 months old animals, the frequency of CD68^+^ microglia isolated from APP/PS1 mice was significantly increased compared to age-matched WT, while the brains of APP/PS1- IL37tg mice did not show significantly increased numbers of CD68^+^ microglial cells (FACS 9-12m F(2, 11)=14.39 p=0.0008; p=0.0008; p=0.0529; p=0.0494) (Fig. 6D-E). The microglial CD11b^+^/CD45_low_ cell population isolated from the brains of 6 and 20-23 months old APP/PS1 and APP/PS1-IL37tg had an increased percentage of CD68^+^ cells compared to those from WT controls (FACS 6m F(2, 12)=13.45 p=0.0009; p=0.0007; p=0.0263; p=0.0985; FACS 20-23m F(2, 12)=13.38 p=0.0009; p=0.0007; p=0.0359; p=0.0383) (Fig. 6F-G). However, APP/PS1-IL37tg had a significantly lower percentage of cells expressing CD68 compared to APP/PS1 mice (Fig. 6F-G).

**Fig. 6.**
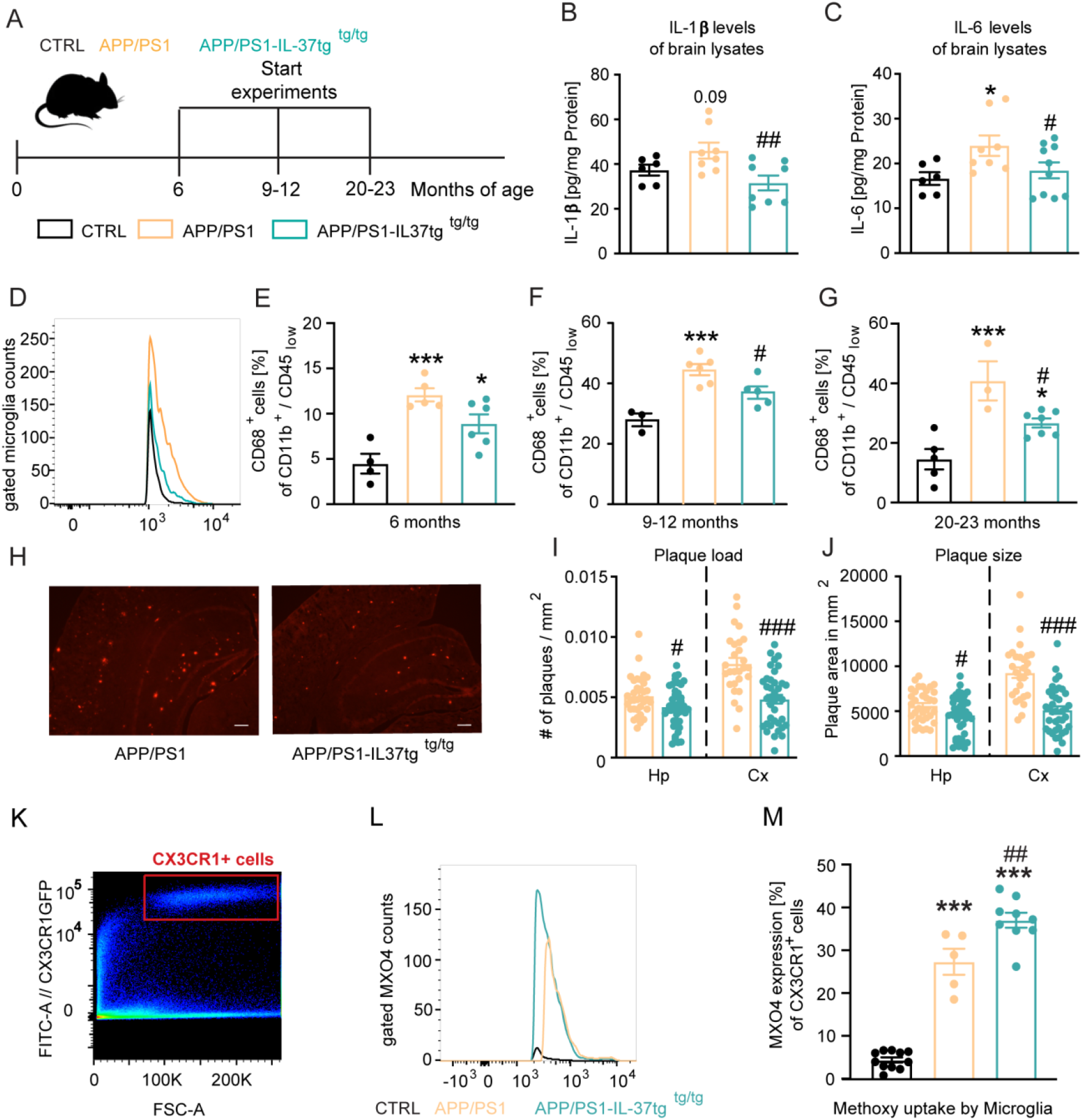
APP/PS1-IL37tg double transgenic mice showed together with less pro-inflammatory cytokine expression and a reduced activation of microglia also fewer number of amyloid-plaque compared to APP/PS1 mice. **(A)** Aged WT, APP/PS1 and APP/PS1-IL37tg mice were analyzed at 6, 9-12 and 20-23 months of age. **(B-C)** Pro-inflammatory cytokine levels of IL-6 and IL-1b were increased in 9-12 months old APP/PS1 mice compared to WT mice, whereas APP/PS1-IL37tg mice showed no enhancement of pro-inflammatory cytokines. **(D-G)** Microglia cells were identified as CD11b+ and CD45low cells and analyzed for their CD68 expression. 9-12 months old APP/PS1 mice showed significant increased percentage of cells with CD68 expression compared to WT mice, whereas APP/PS1-IL37tg mice exhibited no significant enhancement **(D-E)**. Although the 6 and 20-23 months old APP/PS1-IL37tg mice exhibited a significant higher amount of CD68 expressing cells compared to WT mice, the showed a reduction in the percentage of CD68 expressing cells compared to the APP/PS1 animals. **(H-J)** The plaque analysis showed significant lower plaque load **(I)** and reduced plaque size **(J)** in the area of the hippocampus (Hp) and Cortex (Cx) comparing APP/PS1-IL37tg mice to APP/PS1. Representative image of Congo Red staining in 30 µm slices; scale bar 500 µm **(H)**. **(K-M)** Aβ uptake by microglial cells by measuring single cells in the FACS system. Gating strategy for positive Cx3CR1-GFP cells and positive staining for MXO4 **(K-L)**. Quantified analysis of Aβ uptake showing significant higher uptake in APP/PS1-IL37tg cells compared to APP/PS1 cells **(M)**. Data are presented as mean ± SEM. * p < 0.05, ** p < 0.01, *** p < 0.001 compared to WT, # p < 0.05, ## p < 0.01, ### p < 0.001 compared to APP/PS1. (B-G and M: 1-way ANOVA with multiple column comparison; I-J: t-test).

We then examined amyloid-β (Aβ) plaque load in the hippocampus (Fig. 6I) and cortex (Fig. 6J) of APP/PS1 (Fig. 6H left panel) and APP/PS1-IL37tg animals (Fig. 6H right panel). We observed that APP/PS1-IL37tg mice had significantly fewer plaques in both the hippocampus and cortex areas as well as smaller plaque size compared with APP/PS1 mice (plaque burden Hp t=2.403, df=74 p=0.0187; Cx t=4.953, df=63 p<0.0001; plaque size Hp t=2.296, df=74 p=0.0245; Cx t=5.949, df=63 p<0.0001) (Fig. 6H-J). To further analyze whether the observed lower Aβ plaques in APP/PS1-IL37tg were due to higher Aβ clearance, an Aβ uptake assay was performed to analyze phagocytic activity in APP/PS1 and APP/PS1-IL37tg mice carrying the Cx3CR1-GFP gene (heterozygous) (50). For this purpose, methoxy-XO4 staining was performed, followed by FACS analysis. The Cx3Cr1-GFP transgenic mice express a microglia-specific green fluorescent protein in the CNS. To stain Aβ-plaques, animals were injected i.p. with 10 mg/kg methoxy-XO4 3 hours before the start of the experiment. After isolating the brain and performing a single-cell suspension, cells were measured in the FACS system and gated for Cx3CR1-positive cells (Fig. 6K) and then further gated for the methoxy-XO4-positive population. Data showed increased uptake of Aβ-particles by microglial cells from APP/PS1-IL37tg animals compared to those from APP/PS1 mice (F(2, 22)=136.2 p<0.0001; p<0.0001; p<0.0001; p=0.0026) (Fig. 6L-M).

The APP/PS1 mouse model of Alzheimer’s disease shows cognitive deficits as early as 8 months of age (46, 51). To investigate whether IL-37 could affect learning and memory in this animal model, we performed the Morris water maze (MWM) behavioral test on 9- 12 months old control, APP/PS1, and APP/PS1-IL-37tg animals (Fig. 7A). During the 8- day acquisition phase in the MWM, escape latency decreased progressively in all groups (Escape latency F(2, 30)=10.5 p=0.0003) (Fig. 7B). However, APP/PS1 animals showed increased escape latency on day 1 and day 3 to 6 of the training period (Escape latency F(2, 30)=10.5 p=0.0003; WT vs. APP/PS1 day 1 p=0.0463; day 3 p=0.007; day 4 p=0.0017; day5 p=0.0015; day 6 p=0.005) (Fig. 7B). Subsequently, the reference memory test (probe trial) was performed on day 3 before the training and on day 9 24 hours after the last training session. During the probe trials, the mice were tested without the presence of the escape platform. The percentage time that the mice spent in each quadrant was measured, and the target quadrant (TQ) preference was compared with the three non-target quadrants (NT). Control mice showed an explicit preference for the target quadrant on day 9 (t=11.45, df=22 p<0.0001), while in comparison APP/PS1 animals lacked any preference for each of the quadrants (t=0.9874, df=18 p=0.3365) (Fig. 7C). In contrast, APP/PS1-IL37tg mice showed a significantly higher preference for the target quadrant similar to what was observed in control animals (APP/PS1-IL37tg PT t=3.705, df=20 p=0.0014) (PT F(2, 30)=8.415 p=0.0013; p=0.0009; p=0.0505) (Fig. 7C).

**Fig. 7.**
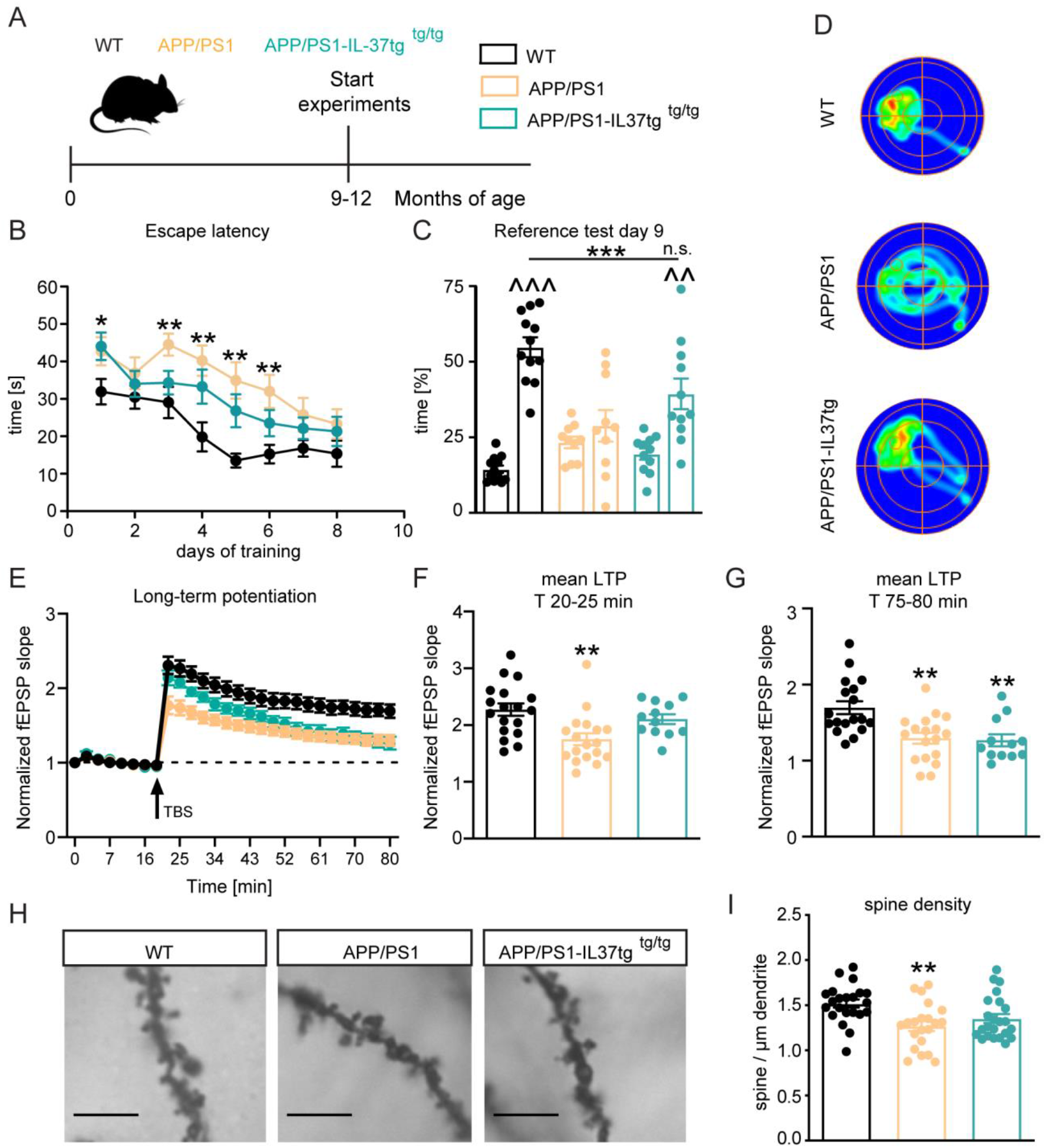
APP/PS1-IL37tg double transgenic mice showed improvements in behavior tests and synaptic plasticity compared to APP/PS1 mice. **(A)** Aged WT, APP/PS1 and APP/PS1- IL37tg mice were analyzed at 9-12 months of age. **(B-D)** The cognitive deficits in APP/PS1 mice in the spatial learning test of the Morris Water Maze could be restored in APP/PS1-IL37tg animals. WT, APP/PS1 and APP/PS1-IL37tg mice indicated a learning behavior during the training phase of the spatial learning test. APP/PS1 animals show higher escape latency during acquisition on day 3-6 compared to WT mice **(B)**. WT mice and APP/PS1-IL37tg mice display a significant preference for the Target Quadrant (TQ), whereas the APP/PS1 mice did not show any preference **(C)**. Representative heat maps of mice of each group manifest the results of the reference test **(D)**. **(E-G)** The LTP deficits in APP/PS1 mice could be rescued in APP/PS1-IL37tg mice in the induction phase (20-25min). However, LTP deficits were not recovered in APP/PS1- IL37tg animals in the maintenance phase (75-80min). **(H-I)** The dendritic spine density was significantly reduced in APP/PS1 animals compared to WT, whereas there was no significant reduction in spine density in APP/PS1-IL37tg. Scale bar 5 µm. Data are presented as mean ± SEM. * p < 0.05, ** p < 0.01, *** p < 0.001 compared to WT, ^^ p < 0.01, ^^^ p < 0.001 compared to NT (non-target quadrants); (B+E: 2-way ANOVA with multiple comparison; C: t-test; C-I: 1-way ANOVA with multiple comparison).

Moreover, heat maps of different groups (1 example per group) were also shown to provide a clearer assessment of the animals’ performance in the reference test. These heat maps show a prolonged time of control and APP/PS1-IL37tg mice in the target quadrant, whereas APP/PS1 mice did not show this behavior (Fig. 7D).

The observed improvement of learning and memory in APP/PS1-IL37 transgenic mice prompted us to analyze whether hippocampal network function could also be improved by this genotype. Therefore, we measured synaptic plasticity in these animals at the Schaffer collateral pathway as described above. Acute hippocampal slices from APP/PS1 mice showed significant deficits in long-term potentiation (LTP) compared to corresponding slices from age-matched control mice (F(2, 45)=9.286 p=0.0004) (Fig. 7E). This was evident during both the induction (20-25min of recording) and maintenance (75-80min of recording) phases of LTP (first 5min F(2, 45)=7.09 p=0.0021; p=0.0016; last 5min F(2, 45)=9.354 p=0.0004; p=0.0014) (Fig. 7F-G). Notably, the induction phase (but not the maintenance phase) of LTP was also indistinguishable in the slices from control and APP/PS1-IL37tg mice (first 5min F(2, 45)=7.09 p=0.0021; p=0.5444; last 5min F(2, 45)=9.354 p=0.0004; p=0.0023) (Fig. 7F-G). Thus, it is clear that expression of IL-37 at 9 months of age can rescue the induction phase of LTP, which may be sufficient for the mice to perform spatial memory tasks.

To determine the rescued phenotypes observed in APP/PS1-IL37tg mice at the cellular level, neuronal morphology of the CA1 hippocampal subregion was analyzed in all experimental groups. A significant reduction in the density of dendritic spines was observed in 9-12 months old APP/PS1 mice compared to age-matched control mice (Spines F(2, 63)=6.318 p=0.0032; p= 0.0026) (Fig. 7H-I). However, a comparable significant difference in the dendritic spine density between control and APP/PS1-IL37tg mice was not observed, indicating a rescue effect by IL-37 (Spines F(2, 63)=6.318 p=0.0032; p= 0.0536) (Fig. 7H-I). Taken together, these results suggest a protective role of IL-37 in transgenic animals by improving the learning and memory deficits associated with the APP/PS1 mouse model and rescuing the underlying cellular correlations.

## Discussion

The regulation of inflammation is an extremely complex and tightly controlled signaling event in which cytokines are crucial modulators of the process (52). The interplay and balance between pro- and anti-inflammatory components are crucial for the delicate balance of inflammatory responses. Anti-inflammatory cytokines, such as IL-1 receptor antagonist (IL-1Ra), IL-10 or IL-37 are known to limit persistent or excessive inflammation, but may also be insufficient or overcompensated (52–54). In this study, we demonstrate for IL-37, an anti-inflammatory cytokine, that it reduces both acute and chronic neuroinflammation and that the IL-37tg mouse is a valuable animal model to target human neurodegenerative diseases caused by chronic inflammation.

The acute inflammatory response was analyzed using a sepsis model induced by systemic administration of the bacterial endotoxin lipopolysaccharide (LPS). Mouse models of septic shock have been shown to have elevated levels of IL-6, TNF-α, and IL- 1β produced by activated microglia (55, 56). The ability of IL-37 to suppress pro-inflammatory cytokines induced by Toll-like receptors (TLR) was first demonstrated in mouse macrophages transfected with human IL-37 (7) and recently described for TLR-induced activation of microglial cells (57). Because, there is no homologues gene for IL- 37 in mice, transgenic mice expressing the human IL-37 gene were generated (IL-37tg) (7, 25). Here we show the effect of IL -37 on CNS tissue resident macrophages in terms of reducing inflammatory markers similar to those in the periphery (Figure 1). The timing of LPS stimulation is important, as longer treatment of 24 hours results in higher cytokine release compared to 6 hours (58, 59). Here we show a significant reduction in pro-inflammatory cytokines from IL-37tg microglial cells after a stimulation period of 24 hours. However, 6 hours with 100ng/ml LPS showed no significant reduction (Fig. 1B). These might be due to the lower stimulus concentration and shorter duration of LPS treatment, which resulted in a weaker response of control microglial cells.

In previous studies IL-37 reverses the metabolic cost of inflammation in plasma and muscle cells by reducing levels of succinate, a potent mediator in inflammatory states (40, 60–62). In addition, macrophage activation has been found to increase production of itaconate, which in turn inhibits succinate dehydrogenase (SDH), causing an increase in succinate levels (37, 63). In this study, primary microglia from IL-37tg animals exhibited reduced levels of itaconate and succinate after LPS stimulation (Figure 2). In summary we demonstrated that the anti-inflammatory property of IL-37 is able to reduce inflammation in microglial cells *in vitro*. Moreover, the restoration of metabolic cost plays a role in rescuing cognitive decline, as shown previously (64).

Further studies were performed by peripheral LPS injection *in vivo* in control and IL-37tg animals. Inflammation and microglial activation are critical hallmarks of various neurological diseases (65). On the one hand, microglial activation is essential for homeostasis in the brain for an adequate inflammatory response. On the other hand, overactivation can lead to neuronal damage (65–67). Systemic inflammation is induced by intraperitoneal injection of LPS. In acute inflammation, little LPS is likely to enter the brain because of an intact blood-brain barrier (BBB) (68). It is more likely that the peripherally-induced neuroinflammation is an indirect effect (65), in which LPS activates the inflammatory cascade that signals to the CNS via TLR and/or TNF-α receptors on the BBB (65, 69–75). Systemic inflammation induces increased density and reactivity of microglial cells characterized by increased secretion of pro-inflammatory cytokines such as IL-1β (56), by changes in the morphological shape (retraction of the processes) (75, 76), by strongly upregulated levels of surface proteins such as CD68 (47, 75, 76), and by increased numbers of IBA-1-positive cells (75). In this study, we show that peripheral induced inflammation leads to significantly increased numbers of IBA-1-positive cells and upregulated levels of CD68 and IL-1β in the CNS of control animals. In contrast, IL-37tg mice display anti-inflammatory effects and did not represent an activated microglial phenotype in these hallmarks (Figure 3). However, the typical morphological changes described as retraction of processes (75) were not observed in either genotypes. IL-37 was originally described as a basic inhibitor of innate immunity with reduced pro-inflammatory cytokine levels in plasma (7) and particularly in peripheral organs and diseases such as endotoxemia, spinal cord injury, colitis, myocardial ischemia, obesity and metabolic syndrome (7, 25, 27–30, 77). Here our findings show *in vivo* neuroinflammatory processes are reduced in acute inflammatory challenged mice with transgenic IL-37 expression.

The resulting neuroinflammation and activation of microglial cells after a peripheral LPS challenge also lead to functional and structural changes in the neuronal network, especially impairment of LTP (44, 78–81) and affects spine morphology with influence on dendritic spine density (82–86). Here, we demonstrate that LTP was impaired in control animals after inflammatory challenge with LPS, whereas IL-37tg animals had no defects in long-term potentiation (Figure 4). In addition, the spine density of CA1 pyramidal neurons and neurons from the dentate gyrus was reduced in control mice after LPS stimulus. The loss of spines was rescued in IL-37tg stimulated with LPS, these mice showed no differences in the number of spines compared to control treated IL-37tg mice (Figure 4). These data represent harmful neuronal changes, which are known to be associated with acute inflammatory processes in the CNS, are reduced in IL-37tg mice. Further electrophysiological experiments with acute slices from control and IL -37tg mice after direct treatment with LPS in the chamber suggested that IL-37 expression directly in the CNS may be responsible for the abrogation of impairments after peripheral stimulation. In addition, we confirmed the mechanisms of the anti-inflammatory action in IL-37tg mice by using the recombinant IL-37 in WT mice. Recombinant human IL-37 is protective against endotoxemia, acute lung and spinal cord injury, asthma and myocardial infarction (11, 25, 28, 35, 36, 87–89). Next, we investigated whether recombinant IL-37 has a similar effect on inflammatory processes in the CNS. In these experiments, our findings show reduced levels of the pro-inflammatory cytokines IL-6 and IL-1α after systemic LPS or IL-1β in wild-type animals pre-treated with rIL-37 compared to the same challenge in wild-type mice (without rIL-37) (Figure 5). The consequences of acute peripheral stimulation by LPS or IL-1β are the induction of pro-inflammatory cytokines leading to impaired learning and memory (90–95). However, IL- 37 expression rescued the impaired memory. The effect of reduced neuroinflammation was also analyzed on cognition and was demonstrated by behavioral improvements in the Y-Maze test of animals pre-treated with rIL-37. Spontaneous alternation behavior (SAB) in a symmetrical Y-shaped maze was used 30 years ago for pharmacological studies of short-term memory performance (96–99). In this first part, we show that acute neuroinflammation, microglial activation, deficits in functional and structural plasticity, and impaired cognition induced by a peripheral stimulus were reduced or rescued in the presence of IL-37 (either as a transgene or recombinant). Furthermore, these results (recombinant IL-37 reduces pro-inflammatory cytokines in the CNS) indicate an important future role for rIL-37 as a pharmaceutical application.

To analyze chronic neuroinflammation, we used APP/PS1 mice as a model for Alzheimer’s disease (AD). AD is the most common form of dementia and in 2015, an estimated 46.8 million people worldwide were living with the dementia (100). Research on AD has mainly been focusing on the long-lasting known pathological hallmarks namely beta-amyloid plaques and neurofibrillary tangles (101–104). Recently, reports linking neuroinflammation to the pathogenic process of AD have been accumulating, demonstrating that the brain can no longer be considered as an absolutely immune-privileged organ in disease progression (101, 104–106). In the present study, we focused on reducing the neuroinflammatory response by decreasing microglial cell activation and the release of pro-inflammatory mediators. For this purpose, we used the IL-37tg mouse line described above. To investigate the role of the anti-inflammatory cytokine IL-37 in chronic inflammation, we crossed the IL-37tg mouse with the APP/PS1 mouse. The APP/PS1 mouse line represents a reliable model that exhibits amyloid-β (Aβ) deposition as early as 6 months of age and is reported to have cognitive deficits beginning at 8 months of age (46, 51, 107). Microglial cells, the resident-immune cells of the brain, are described (along with astrocytes) as a major source of cytokines that have a significant and distinct impact on the neuroinflammation aspects in AD (105, 108). It has been previously described that microglia respond to Aβ peptides, which might be able to trigger the inflammatory process in AD that contributes to microglial activation, release of pro-inflammatory cytokines and memory deficits (104, 105, 109). Here, we show reduced pro-inflammatory cytokine levels for IL-1β and IL-6 in brain lysates from 9 to 12 months old APP/PS1-IL37tg mice compared to APP/PS1 animals. In addition, microglial activation was significantly higher in APP/PS1 mice compared to control mice, whereas APP/PS1-IL37tg mice did not show a significant increase in microglial activation due to CD68 expression. Furthermore, we analyzed the Aβ depositions in the brain tissue of APP/PS1 animals compared with APP/PS1-IL37tg mice. Overexpression of the anti-inflammatory cytokine IL-37 in APP/PS1 animals used in this study resulted in significantly higher plaque phagocytosis capacity, lower plaque burden, and smaller plaque size compared to APP/PS1 control mice. In conclusion, reduction of pro- inflammation or inhibition of excessive inflammation due to IL-37 expression could lead to better Aβ clearance.

It is noteworthy that previous studies have shown that IL-1β and IL-6 play a major role in the progression of AD, as both of them were released from microglial cells surrounding Aβ plaques in AD patients and animal models (104, 110–112). Furthermore, IL-1β and IL-6 are described to play complex roles in regulating cognitive function in AD (95, 104, 113–115). The present results show impaired spatial memory impairment in 9 to 12 months old APP/PS1 mice compared with control animals. However, APP/PS1-IL37tg mice showed partially rescued spatial memory performance compared to APP/PS1 mice. In addition, APP/PS1-IL37tg mice did not exhibit any impairment during the reference memory test.

Given the observed impairment in spatial learning in APP/PS1 animals, we further analyzed synaptic plasticity by measuring LTP as a cellular correlate of learning and memory in each of the three genotypes. These findings demonstrate that the maintenance and induction phases of LTP were reduced in APP/PS1 mice compared to control animals. In contrast, the induction phase of LTP in APP/PS1-IL37tg mice was similar to that in control animals. However, the maintenance phase of LTP was also reduced in APP/PS1-IL37tg animals. These observations indicate that overexpression of IL-37 is sufficient to rescue the early phase of LTP (E-LTP; the first 20 minutes after stimulation), which is independent of protein synthesis. However, impaired late LTP in APP/PS1 mice, which requires regulation of gene expression at the transcriptional and translational levels (116), was not rescued by IL-37 overexpression. Interestingly, there is evidence that the water maze task can be efficiently solved despite late-LTP impairments in the CA1 or the CA3 hippocampal subregion (117, 118). In addition, neuronal morphology was analyzed to investigate whether structural cellular changes are the reason for the reduction in LTP. When APP/PS1 mice were compared with control mice, it was found that APP/PS1 mice had decreased spine density in the apical dendrites of pyramidal CA1 neurons. In contrast, APP/PS1-IL37tg did not show a reduced number of dendritic spines.

In summary, during chronic neuroinflammation associated with Alzheimer’s disease, a significant deficit in synaptic plasticity (functional LTP and structural spine density) was documented, whereas rescue was observed in transgenic AD animals overexpressing the anti-inflammatory cytokine IL-37. Furthermore, a significant effect on the formation of amyloid-β plaques, a pathological hallmark of AD, was documented, suggesting that attenuation of neuroinflammation combined with increased clearance is sufficient to produce beneficial effects on learning and memory. Indeed, the results reported here show that expression of an anti-inflammatory IL-37 cytokine is able to reduce neuroinflammation and cognitive decline in a mouse model of Alzheimer’s disease. Therefore, these data provide the basis for recombinant IL-37 as a therapeutic approach and that the IL-37tg mouse is a novel and important model system to explore therapeutic options while providing mechanistic insights into human neurodegenerative diseases.

## Acknowledgments

This work was in part supported by the DFG (SFB854), and by the Helmholtz-Gemeinschaft, Zukunftsthema “Immunology and Inflammation” (ZT-0027) to MK, and NIH Grant AI-15614 (to CAD). We acknowledge Philip Bufler, for providing us the hIL- 37tg mouse and Andreas Holz for excellent guidance and critical comments to an earlier version of the manuscript.

## Author Contributions

N.L.: performed the experiment, wrote the paper, S.H., M.O.: performed experiments, K.H. contributed reagents; C.A.D: wrote the paper, provided reagents, M.K. designed the experiment, wrote the paper.

## Materials and methods

All experiments were performed and analyzed blinded to the experimenter.

### Animals

All animals used in this study were of either sex. Mice were bred and kept at the animal facility of the TU Braunschweig under standard housing conditions in a 12:12 light:dark cycle at 22°C with food and water available *ad libitum*. Transgenic mouse expressing human IL-37 (hIL-37tg mice, also referred as IL-37tg) were provided by Prof. Dr. Philip Bufler, Medical University of Munich. IL-37tg animals were originally generated by injecting fertilized eggs from C57BL/6 females with the pIRES IL-37 expression plasmid (Nold et al., 2010). As a starting pair for these experiments here, a heterozygous female was mated with a heterozygous male. Resulting negative animals, heterozygous and homozygous IL-37tg animals were identified by PCR. The distinction between heterozygous and homozygous animals could be clearly analyzed based on the positive mRNA quantity. Further C57BL/6J wild-type (WT) mice and APP/PS1ΔE9 (heterozygous breeding) mice were used. The latter mouse line was crossed with the hIL-37 line to create a double-transgenic mouse line (APP/PS1-IL37). In all experiments negative littermates and/or C57BL/6 wild-type mice were used as controls. All experimental procedures were authorized by the animal welfare representative of the TU Braunschweig and the LAVES (Oldenburg, Germany) (33.19-42502-04-16/2170).

### Lipopolysaccharide (LPS) administration

The systemic immune stimulation with lipopolysaccharide (LPS) was performed by intraperitoneal injection. The body weight of the stimulated animals was monitored to determine the appropriate volume of LPS from *Escherichia coli* (*E.coli* O127:B8, Sigma Aldrich L 3129). Animals were either injected with 2x 0.5 mg/kg LPS or 0.9% sodium chloride (NaCl) as a control.

### Administration of recombinant IL-37 and IL-1β

Wild type mice were intraperitoneal injected on 3 consecutive days with 300 ng recombinant IL-37 per mouse following one injection with 60 ng recombinant IL-1β per mouse.

### Cell culture and LPS administration

Neonatal mouse brains (P3 – P5) were used for culture preparation as shown previously (119). Briefly, the meninges were removed and the brain transferred into HBSS 1X on ice. Using a 10 ml pipette the tissue was transferred into a sterile 50 ml conical tube and centrifuged at 2000 rcf for 5 min at 4°C. The re-suspended pellet (in 5 ml fresh HBSS 1X) was applied on a cell strainer (100 µm pores). Again after spinning as before the pellet was re-suspended in 10 ml culture media (DMEM + 10% FCS + 1% Penicillin/Streptomycin) and transferred into a T-75 flask. The mixed culture was incubated in the flask in an incubator at 10% CO_2_ at 37°C for 2-3 weeks. After 3 days incubation the media was replaced 50% with fresh media. In the following every 7 days the media was replaced completely. After 2-3 weeks the culture has reached confluence and the flasks were shaken at 180 rpm for 3 h at 37°C. The media including the microglia cells was collected without disrupting the astrocyte layer on the bottom of the flask and was centrifuged at 3000 rpm for 10 min at room temperature (RT). Microglia cells were plated in 96-well plate with a density of 6x10^4^ cells/well and were treated with different concentrations of LPS for 24h. In the last hour of the treatment, ATP (5 mM) was added to the cells.

### Cytokine measurement of pro-inflammatory IL-6, IL-1β and TNF-α

Enzyme-linked immunosorbent assay (ELISA) was used to quantify cytokines in either brain homogenates or supernatants of treated primary microglia cells. Mice were deeply anesthetized with CO_2_ and killed via decapitation. Brains were isolated and homogenized in STKM buffer (250 mM Sucrose, 50 mM Tris-HCl, 25 mM KCl, 5 mM MgCl_2_) using the GentleMACS (Miltenyi Biotec) program Protein_01. After centrifugation at 4000 g for 5 min at 4°C the supernatant was again centrifuged for 10 min at 13000 g at 4°C. Brain homogenates (1:2) (diluted in 1% BSA solution) were analyzed using R&D systems ELISA Kits.

### IL-37 mRNA measurement

To quantify the expression of IL-37 in primary microglia, cells were treated with 100 ng/mL LPS for 24 h for seven different time periods (5 Min, 10 Min, 20 Min, 40 Min, 1h, 4h and 24h). Total RNA was isolated using peqGOLD TriFast^TM^. cDNA was prepared using BioLabs ProtoScript II First Strand cDNA Synthesis Kit.

For tissue extraction the RNA purification Kit (Macherey-Nagel) was used. cDNA was prepared by using the High Capacity cDNA Reverse Transcription Kit (Thermofisher).

Then, real-time quantitative PCR (qPCR) was performed using BlueProbe qPCR Mix (Biozym) and the following primer pairs: *IL-37*: 5’-GGG AGT TTT GTC TCT ACT GTG AC-3’(forward) and 5’-CCC ACC TGA GCC CTA TAA AAG-3’(reverse); *GAPDH* 5’- GCC TTC CGT GTT CCT ACC-3’(forward) and 5’-CCT CAG TGT AGC CCA AGA TG-3’(reverse). Expression levels of target mRNA was analyzed using the ΔΔCt method and were normalized to the expression level of the house keeping gene GAPDH, which was used as an internal control.

### Immunohistochemistry

To check the amount and morphology of microglial cells in the brain tissue, brains were isolated and fixed in 4% paraformaldehyde (PFA) for 24 hours and then cryoprotected in 30% sucrose solution in phosphate buffered saline (PBS 1x) for 24 hours. Samples were stored in Tissue-Tek® O.C.T.^TM^ compound (A. Hartenstein Laborversand) at -70°C. 30 µm brains sections were cut using the Cryostat. Using the free floating method these slices were washed in 1x PBS and blocked in 1x PBS solution containing 0.2% Triton X- 100, 10% goat serum and 1% BSA for 1 hour at room temperature (RT). Slices were incubated overnight at 4°C with anti-ionized calcium-binding adaptor molecule 1 (IBA-1) (1:1000; rabbit polyclonal, Synaptic System) primary antibody diluted in 1X PBS, 0.2% Triton X-100 and 10% goat serum. Cy^TM^3-conjugated AffiniPure Goat Anti-Rabbit IgG (H+L) (1:500; Jackson ImmunoResearch) was used as secondary antibody diluted in 1X PBS. Sections were washed as before and stained with 4’,6-diamidino-2-phenylindole (DAPI) (SIGMA) followed by cover-slipping with Fluoro-gel with Tris buffer (Electron Microscopy Sciences).

### FACS analysis

Microglia activation was analyzed by measuring the marker CD68 with the FACS method (119). Single cell isolation was performed using the Adult Brain Dissociation Kit (Miltenyi Biotec Order no. 130-107-677) from Miltenyi and the GentleMACS to homogenize the fresh isolated brains. The cells were resuspended in FACS staining buffer (1xPBS + 1% FCS + 0.1% Na-Azide) and plated in V-bottom 96-well plate. Cells were stained for 30min with CD11b-PerCP (1:50), CD45-APC (1:50), CD68-PE (1:50). The flowcytometry was measured using the BD LRS II SORP and analyzed with FlowJo Software.

### Methoxy-XO4 staining

The phagocytic activity regarding Aβ uptake in microglial cells of APP/PS1 and APP/PS1-IL37tg animals was analyzed by FACS. Here, WT, APP/PS1 and APP/PS1-IL37tg animals was used and crossed with the Cx3Cr1-GFP mouse line. The Cx3cr1- GFP transgenic mice express a microglia-specific (or monocytic-specific) green fluorescent protein in the CNS. To visualize Aβ particles animals were injected intraperitoneal with 10 mg/kg methoxy-XO4 (Abcam; ab142818; blood-brain barrier permeable amyloid-β fluorescent marker) in 50% DMSO/ 50% NaCl (0.9%) 3 hours prior to the start of experiments. After the brain was removed, the procedure was as in section “FACS analysis”.

### Y-Maze test

There are several methods to evaluate cognitive function in rodents. Of these, one method is called “Y-Maze” in which a mouse is placed in a maze with three equal arms (hence the term “Y”) each 32 cm from the center. First the mouse is free to explore the first two arms while the third arm is blocked. After the mouse get to know the existence of the two arms during 3 minutes training, the mouse was returned to its cage for 1.5 hours. Then the mouse is placed back to the Y-Maze with all three arms available. The observer records how often the mouse enters an arm. Due to animal’s natural tendency to explore a newly introduced environment it is expected to be expressed by a high frequency of spontaneous alteration performance (SAP). The SAP score is a triplet of three successive different arm visits (ABC, BCA, CAB, BAC, ACB, CBA).

### Morris Water Maze test

The Morris water maze (MWM) test is an assay to analyze spatial memory formation and retention (120). A circular plastic pool (160 cm in diameter and 60 deep) filled up to 30 cm with opaque water (titanium dioxide, Euro OTC Pharma; water temperature 19-20°C) was used including a 10 cm escape platform submerged 1 cm below the water surface and three visual cues on the walls around the pool. Each day the test was performed in the same conditions (dim light and the same time of the day) by the same experimenter blind to all groups. The ANY-maze software (Stoelting, USA) with a camera above the center of the maze was used to track each trial. A pre-training with a visible platform was performed to guarantee the visual and swimming ability in all experimental groups and in addition to get the animal used to the test situation. The pre-training lasted for three consecutive days with two trials each day (maximum of 60 s each) to reach the platform.

The mice were trained in the Morris water maze for 8 days with the invisible platform located in the northwest (NW) quadrant. Each day, animals were placed from randomly starting positions (SW, S, E and NE) for 4 trials in the water with 5 min intervals. The animals had a maximum of 60 s to find the platform otherwise they were guided to the platform and allowed to stay for additional 15 s. The memory retention was measured by performing a reference test. One reference test was performed on day 3 of the training acquisition (prior to the training session) and another reference test 24 hours after the last training day (day 9). The platform was removed during this reference test. The animals were tracked for 45 s (starting position SE).

### Electrophysiological experiments

To study learning and memory processes on cellular level we did electrophysiological recording experiments in CA1 hippocampal neurons as described before (17, 119, 121). Briefly, mice were deeply anesthetized with 100% CO_2_, killed by decapitation following fast brain removal and transfer into ice-cold carbogenated (95% O_2_ and 5% CO_2_) artificial CSG (ACSF) containing 124 mM NaCl, 4.9 mM KCl, 1.2 mM KH_2_PO_4_, 2.0 mM MgSO_4_, 2.0 mM CaCl_2_, 24.6 mM NaHCO_3_ and 10 D-glucose, pH 7.4. Acute hippocampal slices were prepared (400 µm) using a tissue chopper. The hippocampal slices were placed into an interface chamber (Scientific System Design) and incubated at 32°C in a constant flow rate (0.5 ml/min) of carbogenated ACSF for 2h. Afterwards the recordings were started and field excitatory post synaptic potentials (fEPSPs) were measured in the stratum radiatum of the hippocampus CA1 sub-region. The Schaffer collateral pathway was stimulated using two monopolar, lacquer-coated stainless-steel electrodes (5 MΩ; AM Systems). Long-term potentiation (LTP) was induced after 20 min baseline recording by theta-burst stiulation (TBS) including four bursts at 100 Hz repeated 10 times in a 200 ms interval. This stimulation was repeated three times in a 10 s interval. Only healthy sections with a stable baseline were included in the data set. Using the IntraCell software (version 1.5, LIN) the data set was analyzed.

### LPS administration during electrophysiological recordings

To investigate the acute effect of LPS-induced direct immunostimulation and subsequent local IL-37 expression in the CNS, LPS administration was performed during electrophysiological recordings. For this purpose, acute hippocampal slices from adult control and IL-37tg mice were pretreated with LPS (10 µg/ml) 2 hours before and throughout the recording period.

### Golgi-Cox staining

To analyze the morphology of hippocampal neurons, Golgi-Cox staining was performed like previously described (17, 119, 121). Briefly, mice were deeply anesthetized with CO_2_ and sacrificed by decapitation. The brain was incubated in FD rapid Golgi-Cox staining kit according to the manufacturer’s protocol. The tissue was incubated in a mixture of solution A (potassium dichromate and mercuric chloride) and B (potassium chromate) for at least 14 days at RT in the dark. After incubation of A-B the tissue was placed for 1 week in solution C (tissue protection) at RT. In the following, the brain was blocked in 2% agar and 200 µm thick coronal sections were cut using a vibratome (Leica VT 1000 S). The slices were collected on gelatin-coated glass slides and stained with solution D and E before being dehydrated through graded alcohols and mounted using Permount (Thermo Fisher Scientific).

### Congo-Red staining

To examine the Aβ-plaques in APP/PS1 and APP/PS1-IL37 mice plaques were analyzed for the amount and size using the Congo-Red staining. Mice were deeply anesthetized with CO_2_ and sacrificed via decapitation. Brains were isolated and hemispheres were fixed in 4% PFA for 24 hours and then cryoprotected in 30% sucrose solution in phosphate buffered saline (PBS 1x). Hemispheres were stored in Tissue-Tek (Hartenstein Laborversand) at -70°C and cutted using the Leica Cryostat (CM3050 S) in 30µm slices. The sections were transported on gelatin-coated slides and in the following stained with the Congo-Red manufactorer’s protocol using the FD Congo-Red Solution^TM^ kit (FD Neurotechnologies, Inc.) and mounted with Permount (Thermo Fisher Scientific).

### Imaging and image analysis

To analyze the hippocampal neuron morphology, CA1 and dentate gyrus (DG) cells were imaged in the three-dimensions (z-stack thickness of 0.5 µm) using Axioplan 2 imaging microscope (Zeiss) with a 63x (N.A. 1) oil objective equipped with a digital camera (AxioCam MRm, Zeiss). All selected dendrites were analyzed per spine density via number of spines (counted manually using the ImageJ software) per micrometer of dendritic length more than 50-60 µm which were positioned at least 40-50 µm away from the cell soma.

To analyze microglia morphology microscopic images of anti-IBA-1 were taken within the area of cortex and hippocampus. Images were taken in 3D (z-stack thickness 1 µm) using Axioplan 2 imaging microscope (Zeiss) equipped with an ApoTome module (Zeiss) with a 20X objective (NA, 0.8) and digital camera (AxioCam MRm; Zeiss). IBA-1 positive cells were counted with clearly visible nuclei by DAPI staining for microglia density and the processes of IBA-1 positive cells were analyzed to investigate the activation status of these cells by using the ImageJ software (Wane Rasband NIH, USA).

To survey the Aβ-plaques images from Congo-Red stained slices were taken. Congo-Red staining presents a bright fluorescence emission at 614 nm with excitation at 497 nm. Images of brain sections were taken using an Axioplan 2 imaging microscope (Zeiss) with a 2.5X objective (N.A. 0.07) connected to a digital camera (Nikon) with the same light exposure time of 1 s in all groups. Plaque load and plaque size were analyzed using the ImageJ software (Wane Rasband NIH, USA) with the analyze particle tool. The polygon selection tool was used to generate the region of interest (ROI) for the area of hippocampus and cortex. Plaque load (number of particles) and plaque size (area of particles) were normalized to the area of hippocampus and cortex and plotted as plaque load and plaque size per mm^2^.

### Extraction of intracellular Metabolites for Gas Chromatography-Mass Spectrometry (GC-MS)

Primary microglia cells (5*10^5^ cells) were plated onto 12-well plates and incubated for 24 h. The medium was exchanged with fresh medium or medium mixed with 10 ng/ml LPS for 24 h. To extract intracellular metabolites cells were washed once with 0.9% NaCl and 500 µL of a cold methanol/water mixture was added. The water fraction contained the internal standard (IS) D6-glutaric-acid (c = 1 µg/ml). Cells were scraped and transferred into 250µL of -20°C Chloroform. After <u>vortexing</u> samples for 20 min at 4°C with maximal speed in an automatic shaker, samples were centrifuge for 5 min at above 15.000 xg in a table centrifuge (4°C). 200 µL of the upper phase was transferred in a GC glass vial with micro insert and evaporated to dry under vacuum at 4°C, overnight. To avoid condensation, the GC glass vials were warmed to room temperature under vacuum and capped afterwards with magnetic caps.

### Metabolite Measurement

Derivatization of the samples was performed directly before GC-MS measurement. Metabolite extracts were dissolved in 15 µL pyridine, containing 20 mg/mL methoxyamine hydrochloride at 55°C for 90 min under shaking. After adding 15 µL N-methyl-N-tert-butyldimethylsilyl-trifluoroacetamide samples were incubated at 55°C for 60 min under continuous shaking. GC/MS analysis was performed using an Agilent 7890B GC coupled to an Agilent 5977B MSD. The gas chromatogram was equipped with a 30-m ZB-35 Phenomenex 5-m Guard capillary column. As carrier gas helium was used at a flow rate of 1.0 ml/min. A sample volume of 1 µL was injected into a split/splitless inlet, operating a splitless mode at 270°C. The GC oven Temperature was held at 100°C for 2 min and increased to 300 at 10°C/min and held for further 4 min. Afterwards the temperature was increased to 325°C. The MSD was operating under electron ionization at 70 eV. The MS source was held at 230°C and the quadrupole at 150°C. The total run time of one sample was 26 min. Full scan mass spectra were acquired from m/z 70 to m/z 800. All GC-MS chromatograms were processed using Metabolite Detector, v3.020151231Ra (122).

### Statistical analysis

Data were analyzed and plotted by GraphPad Prism 8 (GraphPad Software, Inc. USA) and presented as mean±SEM. Statistically analysis were performed with either unpaired t-test, one-way ANOVA (post hoc test Fisher’s LSD or Turkey’s multiple comparisons) or two-way ANOVA (post hoc test Fisher’s LSD or Bonferroni’s multiple comparisons) depending on experiments. The minimum significance value was considered as p < 0.05.

## Reagent and resources table

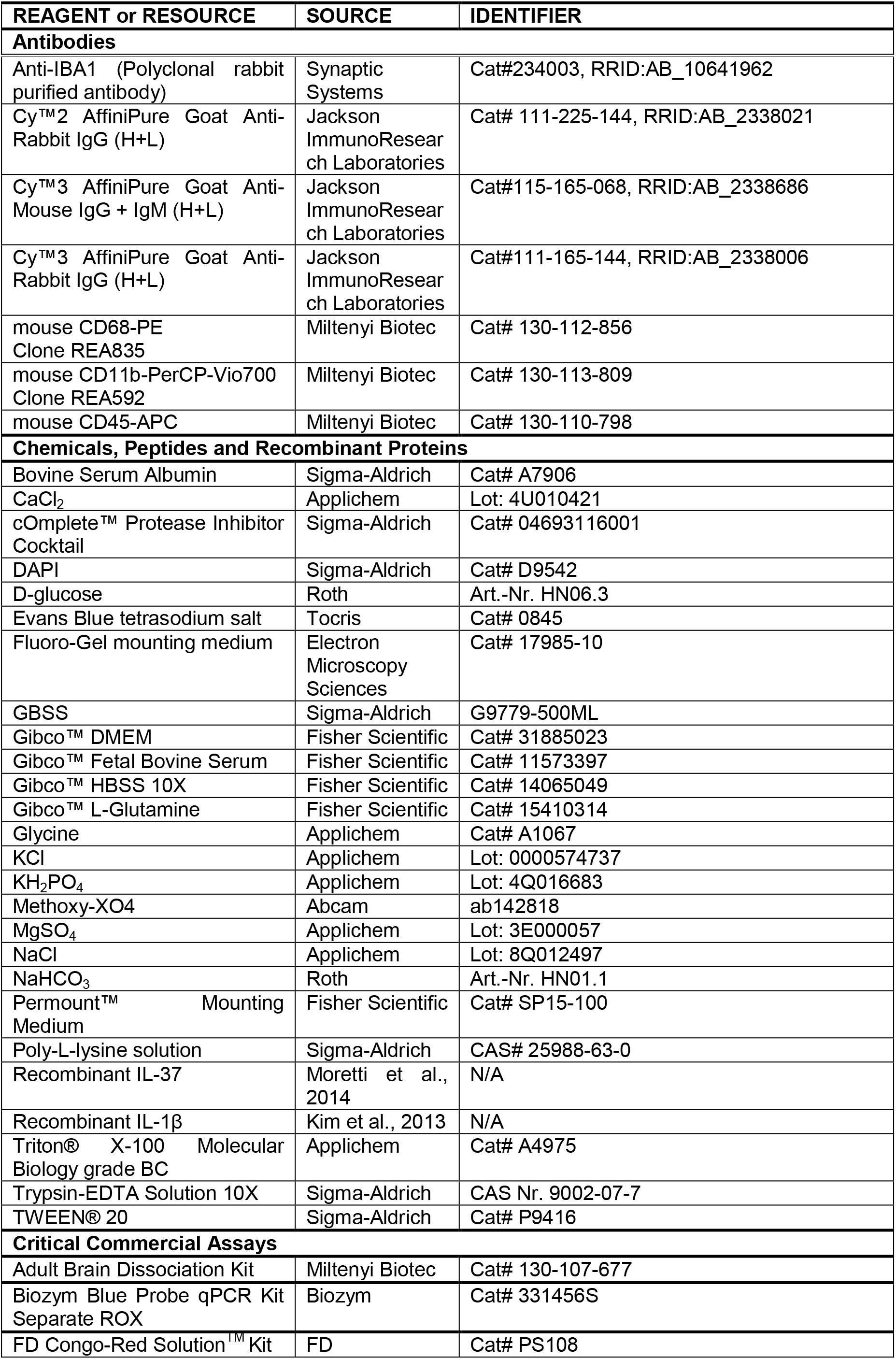
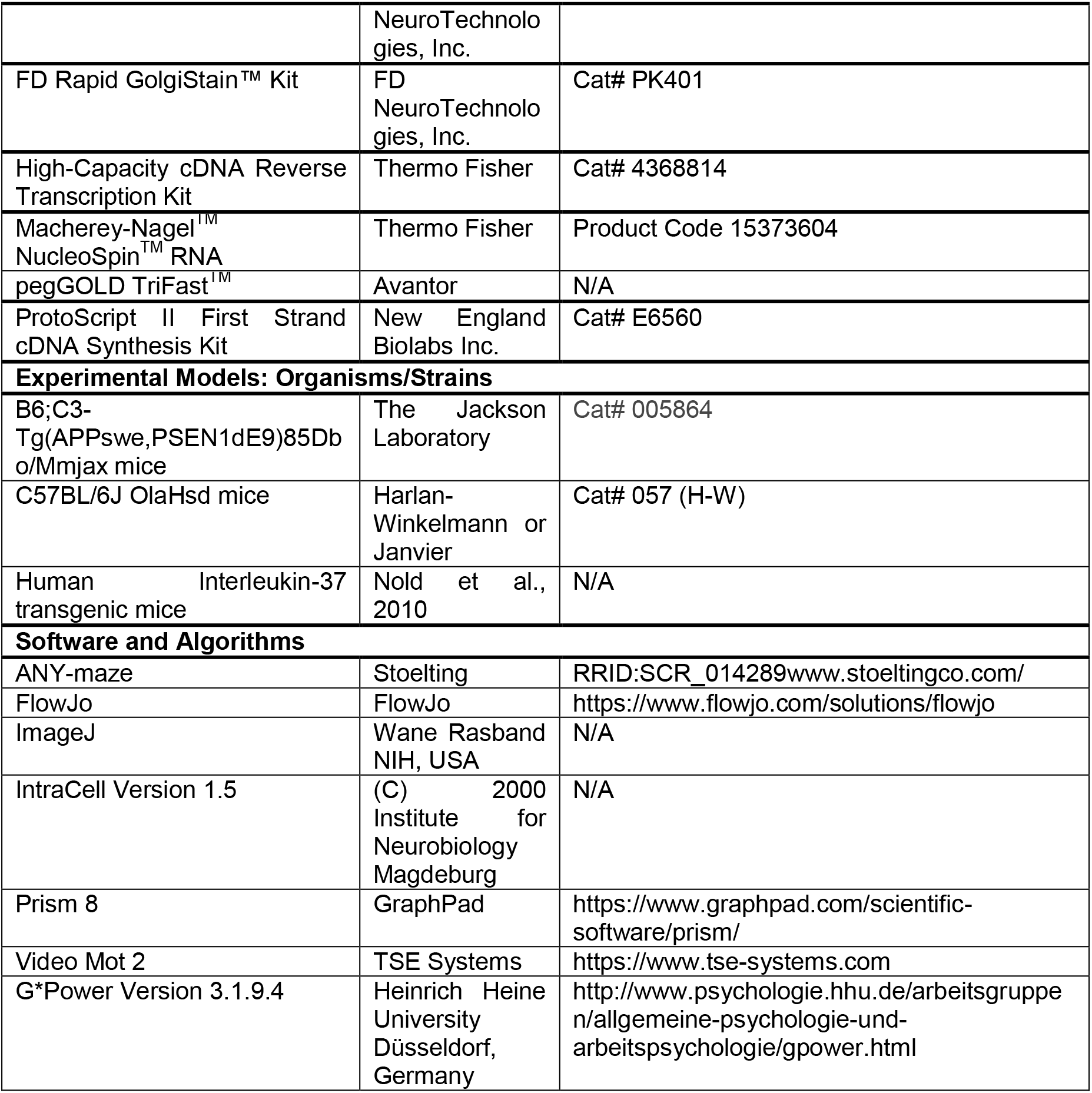

